# Conformational gating of product release sets the catalytic ceiling of a bioluminescent reporter

**DOI:** 10.1101/2025.09.16.675553

**Authors:** M. Toul, J. Horackova, A. Schenkmayerova, J. Planas, T. Landolt, J. Sucharitakul, Y.L. Janin, K. Prakinee, P. Chaiyen, S. Stavrakis, A. deMello, K.A. Johnson, J Damborsky, M. Marek, D. Bednar, Z. Prokop

**Author notes:** These authors contributed equally to this work (M. Toul, J. Horackova).

## Abstract

Luciferases are widely used bioluminescent reporters, yet the molecular determinants of their catalytic efficiency and light-emission stability remain incompletely understood. Here, we reconstruct the complete catalytic pathway of *Renilla* luciferase by combining steady-state, transient, and temperature-dependent kinetics with crystallography and molecular simulations. We show that the enzyme is substantially undersaturated with oxygen 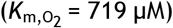, causing its true turnover number (*k*_cat_ = 21.9 s^−1^) to be systematically underestimated. Concurrently, elevated oxygen drives irreversible enzyme inactivation after ~1,500 turnovers, revealing a fundamental trade-off that limits oxygen-affinity engineering. Instead, the genuine bottleneck of the catalytic cycle is the induced-fit conformational opening of the product-bound enzyme. Selective engineering of this transition step through rational loop grafting yielded a variant AncFT-L14 with enhanced catalytic efficiency and glow-type bioluminescence with substantially slower signal decay in cell lysates. Collectively, our results identify conformational dynamics as a primary tunable determinant of luciferase function. More broadly, this work establishes a mechanistically grounded framework for the development of next-generation bioluminescent tools and for engineering enzymes controlled by dynamically gated ligand exchange.

**One-sentence summary:** The slowest step in the widely used *Renilla* luciferase is the conformational opening of the product-bound state and its engineering makes the enzyme glow for longer.

## Introduction

Bioluminescence, a fascinating process of light emission by living organisms, plays crucial roles in nature, including prey or mate attraction, defense, camouflage, and communication(*1–3*). Moreover, bioluminescence has become a cornerstone of modern biotechnology, enabling applications in diagnostics, environmental monitoring, high-throughput drug screening, and gene expression and cellular interaction analyses(*4–6*). Advances in deciphering, harnessing, and engineering bioluminescent systems continue to drive innovation across diverse fields.

Light emission is accomplished by luciferases, enzymes genetically encoded by the corresponding bioluminescent organisms. Luciferases catalyze the oxidative conversion of small-molecule luciferin substrates into electronically excited oxyluciferin products that emit a photon upon relaxation. Despite sharing this general reaction principle, multiple luciferases have evolved independently, are structurally diverse, and use distinct luciferins, overall producing a wide range of emission wavelengths (**Figure 1a**)(*5, 7, 8*). Luciferase from the sea pansy *Renilla reniformis* (RLuc) is among the most popular bioluminescence-based reporters due to its monomeric form, small size of 36 kDa, and cofactor independence. RLuc catalyzes the oxidative decarboxylation of coelenterazine (CTZ) to coelenteramide (CEI), emitting blue light at ~480 nm (**Figure 1b**)(*9, 10*). In 2006, Loening and coauthors reported an 8-point mutant variant of RLuc (RLuc8) with enhanced light output and serum inactivation resistance(*11*). These qualities have made RLuc8 a benchmark standard for many follow-up studies and RLuc-based assay development. Nevertheless, the utility of RLuc8 remains limited by its suboptimal emission spectrum, substantial product inhibition, and, in particular, short luminescence half-life with rapid decay after the initial flash.

**Figure 1:**
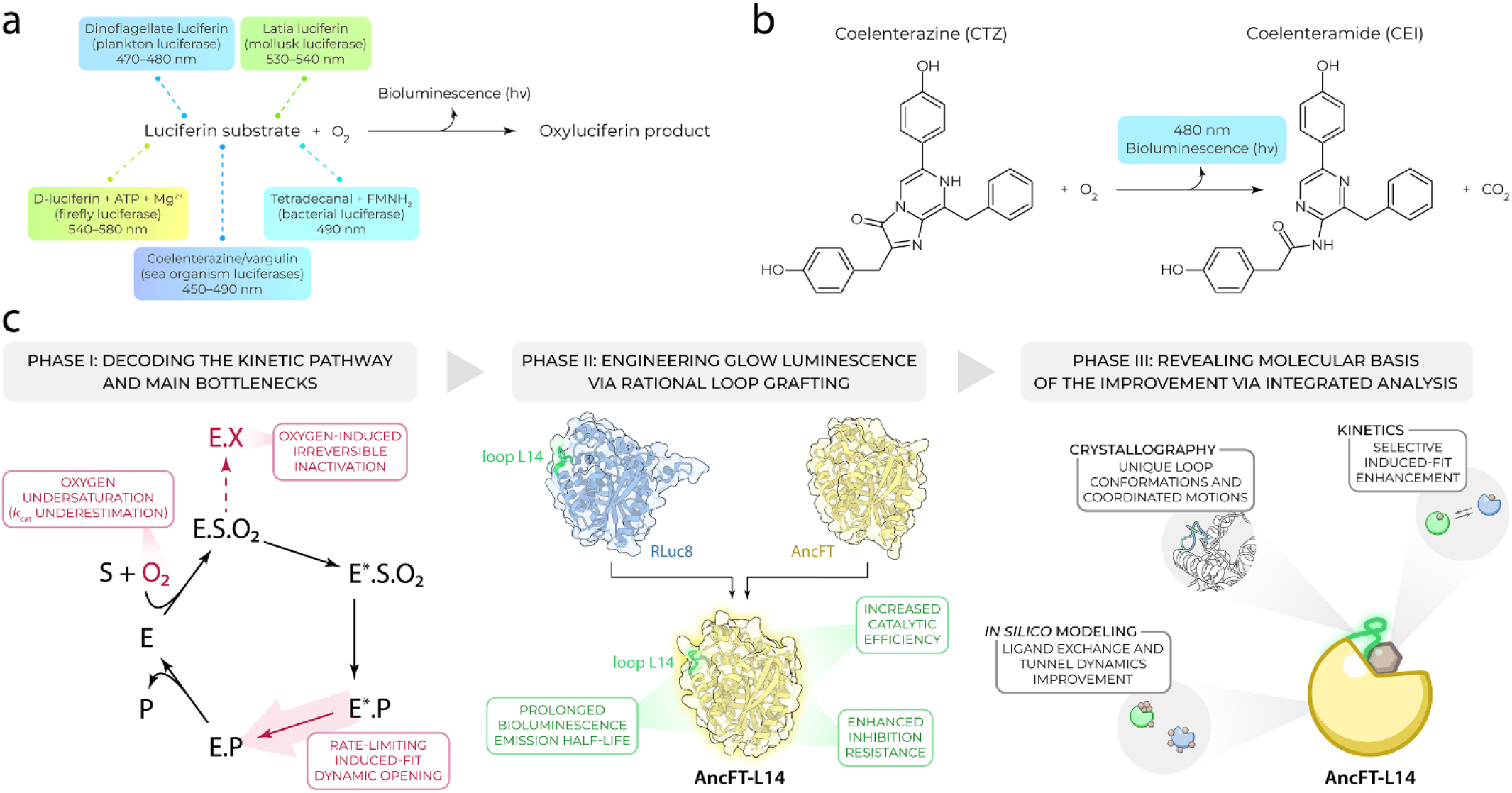
Overview of the luciferase-mediated bioluminescence principle and the mechanism-guided engineering strategy. (**a**) The general scheme common to all luciferases includes the oxidation of a substrate (luciferin) to form a product (oxyluciferin) and emit visible light. Selected examples of specific substrate names and emission wavelengths for individual luciferase types are provided above and below the reaction. (**b**) *Renilla* luciferase-catalyzed oxidative decarboxylation of coelenterazine (CTZ) into coelenteramide (CEI), carbon dioxide (CO_2_), and visible light emission with a maximum at ~480 nm. The colored background corresponds to the color of the emitted light. (**c**) Overall summary and workflow of the work done in this study, including (i) deciphering a detailed kinetic pathway of RLuc8 catalysis with the main bottlenecks, (ii) eliminating the identified limitations by rational loop grafting to generate a luciferase with improved catalytic and light-emission properties, and (iii) integrative kinetic, structural, and computational analysis to rationalize molecular basis of the achieved improvement and to provide a blueprint for follow-up luciferase engineering. E and E* = enzyme in two different conformations; S = substrate; P = product; O_2_ = oxygen; E.X = dead-end inactivated enzyme form.

Our recent work demonstrated that the dynamic behavior of RLuc8 is essential for catalysis. This finding enabled us to generate a luciferase variant with a 100-fold longer light emission half-life compared to that of RLuc8(*12*). This ‘glow-type’ variant, termed AncFT, was produced by transplanting a crucial dynamic fragment from RLuc8 to the reconstructed ancestral luciferase Anc^HLD-RLuc^ that was reported by Chaloupkova *et al*. in 2019(*13*). Furthermore, we recently revealed the catalytic mechanism of RLuc8, identifying key catalytic residues and individual chemical steps and intermediates of this oxidative decarboxylation reaction(*14*). However, crucial steps preceding and following this chemical transformation, i.e., substrate binding and product release, remain poorly understood, leaving the full enzymatic cycle incompletely characterized. These ligand transport steps are often governed by conformational rearrangements of dynamic regions, whose significance in RLuc8 has been previously identified, yet their kinetic contributions have not been resolved. This gap represents one of the last missing puzzle pieces to fully decipher the entire RLuc8 enzymatic pathway and a major barrier to rationally engineer luciferases with tailored light-emission properties.

In this study, we combine steady-state, pre-steady-state, temperature-dependent, and oxygen-dependent transient kinetics to elucidate the detailed kinetic pathway of RLuc8, from initial substrate binding to final product release, and to leverage this mechanistic insight for enzyme engineering (**Figure 1c**). We uncover previously unrecognized oxygen undersaturation, an oxygen-induced irreversible inactivation, and a rate-limiting induced-fit conformational transition associated with product release. By rationally targeting the dynamic conformational bottleneck through loop grafting, we engineer a new variant, AncFT-L14, with substantially accelerated induced-fit dynamics, enhanced catalytic efficiency, and a strongly stabilized glow-type luminescence. These industrially attractive parameters are superior to previously reported *Renilla*-type luciferases(*11–16*). Follow-up kinetic, structural, and computational analyses reveal that coordinated interactions between grafted flexible regions reshape the conformational energy landscape and improve ligand exchange kinetics, providing a molecular basis of the achieved improvement.

Altogether, our findings resolve one of the final mechanistic gaps in the bioluminescence pathway of *Renilla*-type luciferases and establish a mechanistically guided framework for engineering next-generation luciferases with improved emission stability, efficiency, and suitability for quantitative and long-timescale bioluminescence applications.

## Results

### Oxygen-dependent kinetics reveal luciferase irreversible inactivation and turnover number underestimation

Understanding how molecular oxygen influences luciferase catalysis is essential, yet kinetic studies reported to date have varied only the luciferin substrate concentration and assumed the oxygen co-substrate to be non-limiting(*11–18*). To directly examine the contribution of oxygen, we systematically collected two-substrate bioluminescence kinetics by varying both CTZ and oxygen substrate concentrations, and performed a global numerical fitting to an extended steady-state model (**Figure 2a–e**).

**Figure 2:**
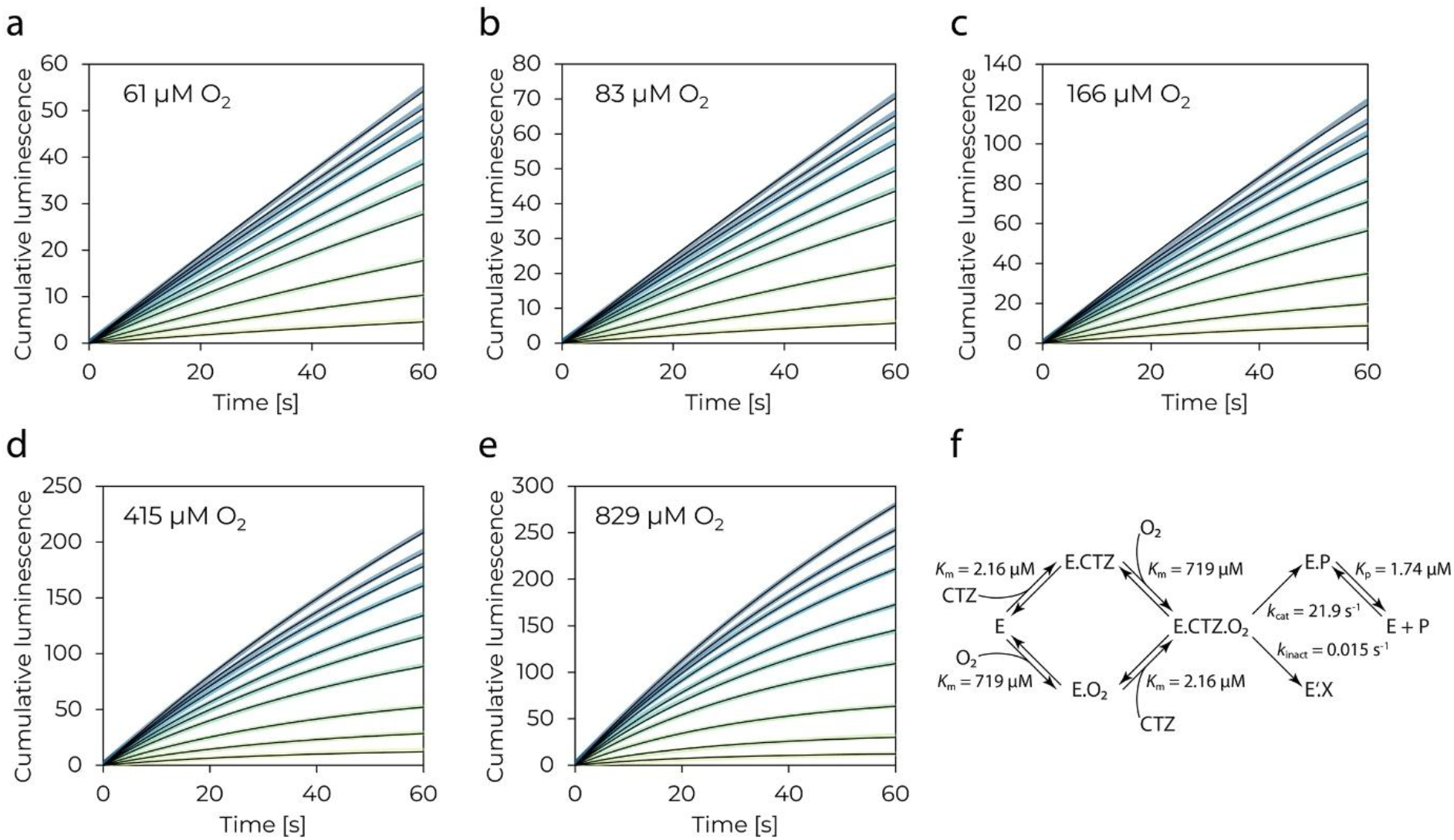
Two-substrate oxygen-dependent steady-state kinetic analysis of the *Renilla* luciferase RLuc8. (**a– e**) Cumulative luminescence progress curves upon mixing RLuc8 with coelenterazine (CTZ) at varying concentrations and oxygen (O_2_) at fixed concentrations. The resulting concentrations after mixing were 0.2–15 µM CTZ (yellow to dark blue curves) and 8 nM RLuc8. The resulting O_2_ concentration is shown for panels (a)–(e) in the top left corner of the graph. All the experiments were performed in 100 mM potassium phosphate buffer (pH 7.5) at 37 °C. The data are shown as averages of 6 technical replicates, and the black solid lines represent the best fit according to the kinetic pathway represented in (f). (**f**) Extended steady-state kinetic pathway for independent binding of CTZ and O_2_ and their conversion by the RLuc8 enzyme (E), leading to the final product (P) or the enzyme-inactive state (E’.X). The analysis allowed the determination of Michaelis constants (*K*_m_) for both substrates, turnover number (*k*_cat_), enzyme inactivation rate constant (*k*_inact_), and product inhibition equilibrium constant (*K*_p_).

The resulting parameters *k*_cat_, *K*_m,CTZ_, and *K*_p_ (**Figure 2f, Figure S1, Table SI**) were consistent with previously reported values(*11–15*), but the analysis further revealed independent binding of CTZ and oxygen and discrete substrate-binding sites in RLuc8. Strikingly, the Michaelis constant for oxygen 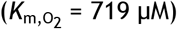 far exceeded the oxygen concentration in air-equilibrated buffer (~210 µM at 37 °C) and is one to two orders of magnitude above intracellular oxygen levels (~5–50 µM), where these reporters are actually used. This finding indicates that RLuc8 is substantially undersaturated with oxygen during catalysis, and severely so in cells. As a consequence, the true turnover number of the enzyme is markedly higher than previously considered: we determined a *k*_cat_ of 22 s^−1^, approximately 5-fold greater than the earlier apparent *k*_cat,app_ values measured under non-saturating oxygen conditions as reported until now (**Table SI**)(*11, 12, 14*). This outcome addresses a longstanding discrepancy in RLuc enzymology and demonstrates that oxygen availability, rather than CTZ concentration, is the primary limitation of the luciferase catalytic performance, pinpointing a potential protein engineering hotspot.

Unexpectedly, the extended global analysis additionally detected a slow but irreversible enzyme inactivation that becomes prominent at elevated oxygen concentrations (**Figure 2f, Figure S1, Table SI**). This behavior is consistent with previous reports of complete activity loss in other CTZ-utilizing luciferases after a defined number of catalytic cycles(*19–22*). Our findings now reveal that RLuc8 is similarly susceptible when exposed to high oxygen levels, suggesting an oxygen-driven inactivation mechanism, likely due to the presence of reactive oxygen radical intermediates, recently proven to be involved in the CTZ oxidation pathway(*14*). The partition ratio between productive turnover and inactivation (*k*_cat_/*k*_inact_ ≈ 1,500) sets a finite catalytic lifetime of roughly 1.5×10^3^ turnovers per enzyme at saturating oxygen. By linking this finite turnover capacity to oxygen radicals via these observations, we provide a deeper mechanistic understanding of this unusual, previously reported inactivation phenomenon.

Above all, these results collectively expose a fundamental trade-off in luciferase engineering. Increasing oxygen affinity by lowering 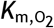 would improve substrate saturation and boost catalytic efficiency by up to 5-fold, but would simultaneously accelerate oxygen-dependent inactivation, ultimately reducing enzyme lifetime and light-emission stability. Thus, while our refined steady-state analysis provided important new mechanistic insights, it also indicated that the challenges in engineering RLuc8 extend beyond oxygen binding. Consequently, targeting oxygen affinity alone would be a suboptimal strategy for further optimization. This outcome prompted a deeper kinetic investigation to identify which step in the catalytic cycle is intrinsically rate-limiting and amenable to protein engineering.

### Transient kinetics pinpoint the rate-limiting induced-fit conformational change as a target for engineering

To achieve a more detailed understanding of the kinetic pathway and identify the rate-limiting step of the RLuc8 catalytic cycle, we complemented our steady-state analysis with a comprehensive pre-steady-state investigation. This work comprised 28 kinetic datasets, including anaerobic substrate binding, product binding, time-resolved spectral measurements, and both oxygen- and temperature-dependent transient kinetic experiments (**Figure 3**). The data were acquired by conventional stopped-flow and quench-flow devices, alongside a droplet-based continuous-flow microfluidic platform previously reported by us(*23*). This microfluidic platform enabled high-throughput data collection with rapid heat transfer (temperature jump), making it particularly suitable for the systematic collection of both transient kinetic and thermodynamic data.

**Figure 3:**
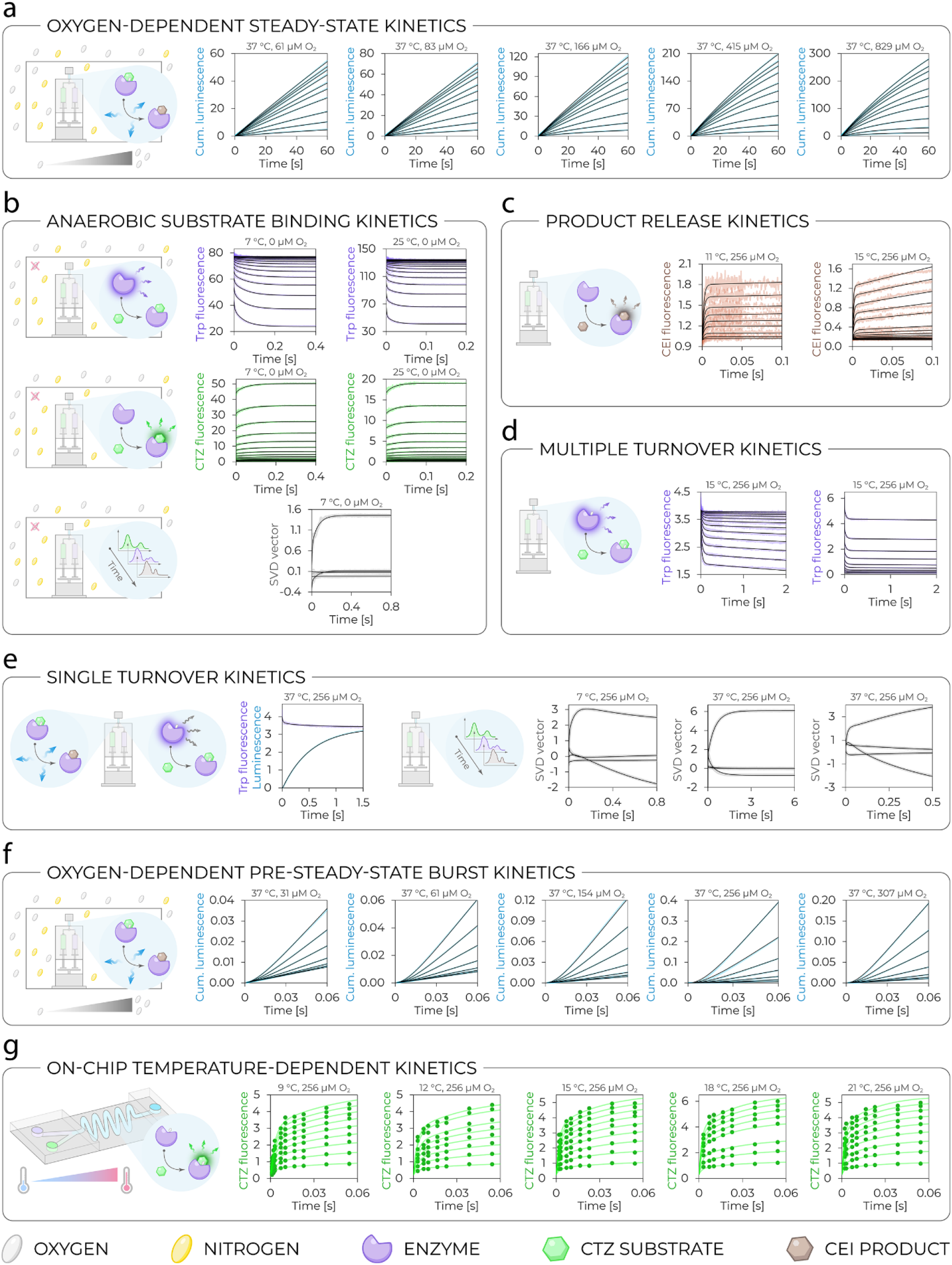
Pre-steady-state transient kinetic data collected for the *Renilla* luciferase RLuc8 and their global numerical analysis. A collection of 28 datasets combines oxygen-dependent steady-state kinetics (**a**), anaerobic substrate binding kinetics (**b**), product release kinetics (**c**), multiple turnover kinetics (**d**), single turnover kinetics (**e**), oxygen-dependent burst kinetics (**f**), and on-chip temperature-dependent kinetics (**g**). The color of the datapoints corresponds to the measured signal, as shown on the Y-axis and illustrated in the adjacent pictograms together with the experimental setup. The black solid lines represent the best fit from the global analysis, yielding the extended kinetic pathway and rate/equilibrium constants displayed in **Figure 4**. The experimental conditions for each individual experiment (concentrations, temperature, buffer, replicates, etc.) are provided in the Online Methods section. The temperatures and oxygen concentrations are also stated above each dataset. CTZ = coelenterazine; CEI = coelenteramide; Trp = tryptophan; SVD = singular value decomposition.

Global numerical analysis of these combined datasets(*24, 25*) allowed us to derive an extended kinetic pathway, revealing several key mechanistic features of RLuc8 catalysis (**Figure 4, Figure S2, Table SII, Supplementary Note 1**). First, the oxygen co-substrate can bind to any enzyme state, i.e., to both the free enzyme (E) and the CTZ-bound enzyme, in either the open conformation (E.CTZ) or the closed conformation (E^*^.CTZ) state. Second, oxygen association is independent of CTZ substrate binding and insensitive to enzyme conformation because the oxygen affinities for all the enzyme states (E, E.CTZ, and E^*^.CTZ) are equivalent. Third, both the CTZ substrate binding and the CEI product release proceed through an induced-fit mechanism in which an initial collision complex undergoes a conformational transition. This behavior underscores the central role of protein dynamics in catalysis, consistent with previous findings(*12*).

**Figure 4:**
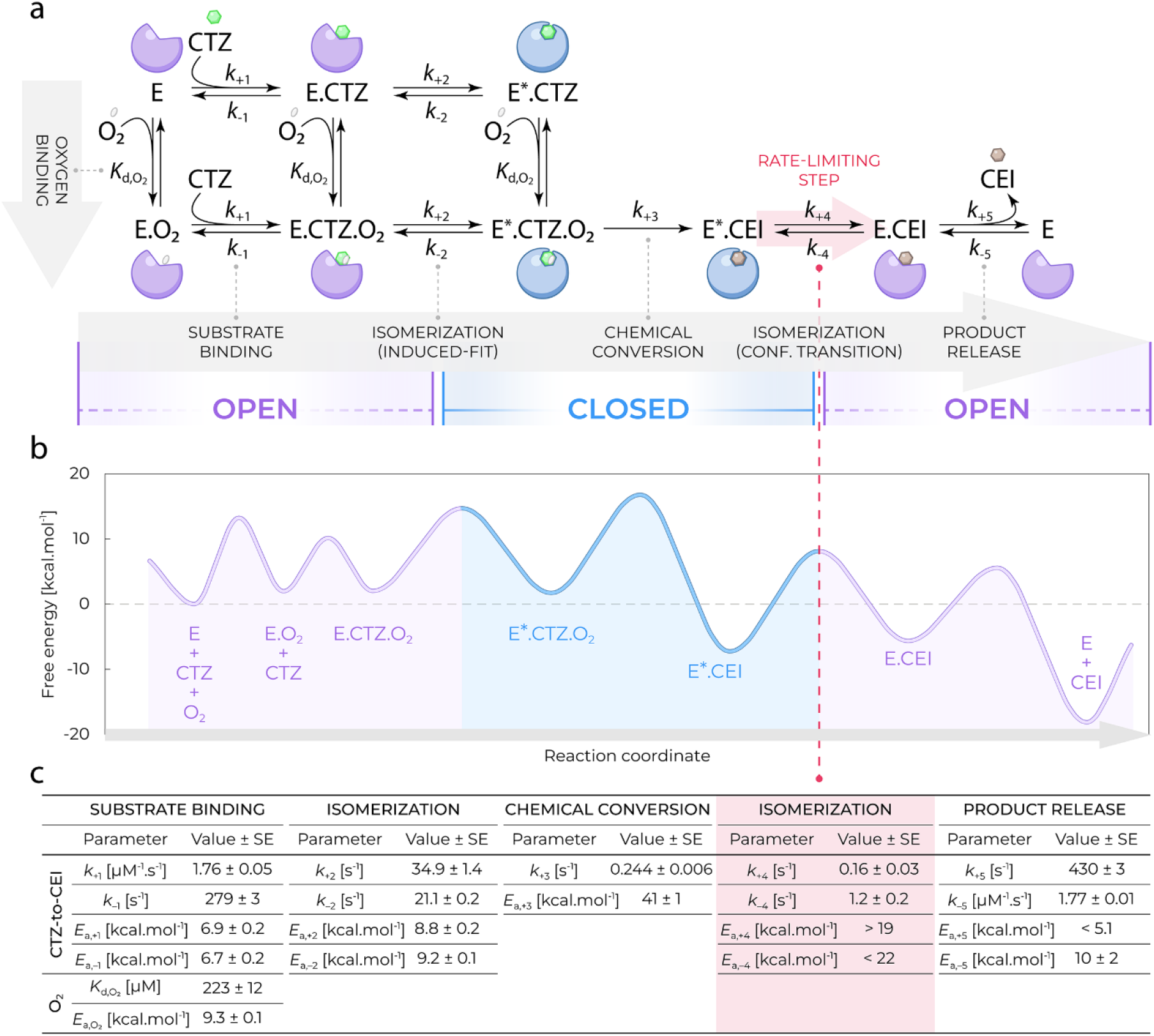
Extended kinetic mechanism of the *Renilla* luciferase RLuc8. Key conformational changes during catalysis, which allow the enzyme to switch between the open and closed states, are highlighted in purple and blue, respectively. The rate-limiting step, a conformational change of the product-bound state, is highlighted in red. (**a**) Graphical illustration of the newly revealed enzyme (E; purple/blue blob) kinetic pathway describing the conversion of the substrate coelenterazine (CTZ; green hexagon) and oxygen (O_2_; gray oval) into the product coelenteramide (CEI; brown hexagon). The pathway includes independent binding of the substrates and critical conformational changes that occur during substrate binding and product release processes. Note that the carbon dioxide product is not included in the pathway as it was not monitored in any of the experiments and thus, no kinetic information is available. E = RLuc8 enzyme; CTZ = coelenterazine substrate; O_2_ = oxygen co-substrate; CEI = coelenteramide product; E.CTZ and E^*^.CTZ = enzyme–coelenterazine complex in two different conformations; E.O_2_ = enzyme–oxygen complex; E.CTZ.O_2_ and E^*^.CTZ.O_2_ = ternary enzyme–coelenterazine– oxygen complex in two different conformations; E.CEI and E^*^.CEI = enzyme–coelenteramide complex in two different conformations. (**b**) Reaction coordinate illustrating the energy barriers of transitions between individual steps of the newly extended kinetic pathway, including the rate-limiting step marked with the red dashed arrow. The free energy profile was derived from the determined rate constants stated in (c) using the transition state theory. (**c**) Values and standard errors of individual rate and equilibrium constants, and activation energies of the newly extended kinetic pathway. The notion of the parameters is per those provided in (a). The reported values correspond to the reaction kinetics at 15 °C in 100 mM potassium phosphate buffer (pH 7.5).

Most importantly, the detailed kinetic analysis further clarified the relative rates of the elementary steps and pinpointed the limiting processes. While the formation and dissociation of the initial CTZ and CEI encounter complexes (*k*_±1_ and *k*_±5_, respectively) are fast, the associated coupled conformational transitions, i.e., enzyme opening and closing (*k*_±2_ and *k*_±4_), are markedly slower. Although the chemical substrate-to-product conversion step (*k*_+3_) is also very slow, it does not represent the major kinetic bottleneck under the assay conditions. Instead, the rate-limiting step is the subsequent induced-fit conformational opening of the product-bound complex, which controls CEI release (*k*_+4_). This single conformational transition imposes the largest barrier in the catalytic cycle and ultimately dictates the overall turnover rate under the conditions of the transient kinetic analysis.

By defining the extended catalytic pathway of RLuc8 and quantifying rate constants of the corresponding elementary steps, our global transient kinetic analysis resolves the key mechanistic gaps in ligand trafficking that remained unanswered after the chemical mechanism was previously uncovered(*14*). Most critically, the identification of the rate-limiting conformational transition provides a clear and selective engineering target, avoiding the inherent trade-offs associated with modifying oxygen affinity revealed by our steady-state analysis. Consequently, these mechanistic insights directly motivated us to focus on modulating protein conformational dynamics, rather than substrate chemistry or oxygen binding, as the principal leverage point for enhancing the overall luciferase catalytic performance.

### Rational loop grafting improves catalytic efficiency and bioluminescence half-life

To test whether the alteration in protein dynamics could selectively accelerate the identified conformational bottleneck and enhance luciferase performance, we implemented a rational loop-grafting strategy. Our previous work demonstrated that flexible active-site loops govern extensive dynamics in *Renilla*-type luciferases(*12*), making their grafting ideal for modulating induced-fit conformational transitions without significantly affecting the catalytic chemistry or initial encounter binding. Using the LoopGrafter web tool(*26, 27*), which integrates loop geometries, lengths, dynamics, and correlated motions, we systematically explored loop exchanges between the herein characterized luciferase RLuc8 and the reconstructed ancestral variants Anc^HLD-RLuc^ and AncFT, previously reported by us(*12, 13*) (**Figure 5a**). LoopGrafter automatically identified correlated motions of loop L9 and loop L14, indicating the potential synergistic benefit of combining both loops within a single scaffold.

Grafting the loop L14 from RLuc8 into AncFT, already containing the transplanted L9 segment, yielded the engineered variant AncFT-L14 with remarkably improved properties (**Figure 5b-f**). Unimpaired circular dichroism spectra confirmed that the grafting did not perturb overall protein folding (**Figure 5b**) and similarly, the emission spectrum of AncFT-L14 remained comparable to that of AncFT (**Figure 5c**). AncFT-L14 further retained a higher melting temperature than RLuc8 (**Figure 5d, Figure S3**), indicating preserved-enough thermostability. At the same time, the L14 graft cost ~10 °C of the melting temperature relative to the AncFT parent, an expected consequence of increasing local loop flexibility and a stability–activity trade-off.

**Figure 5:**
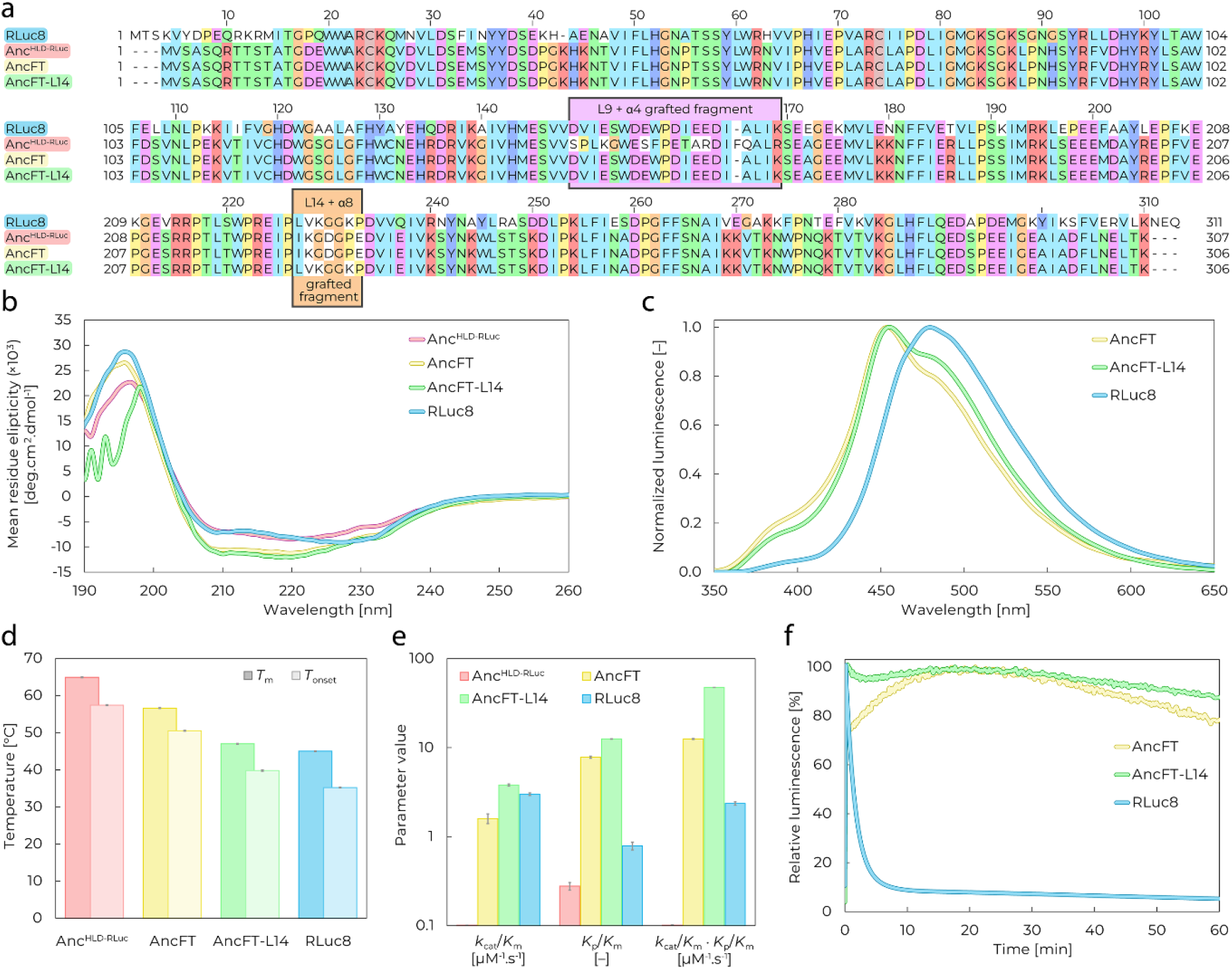
Sequence alignment and experimental characterization of selected luciferases. The comparison includes an 8-point mutant of modern *Renilla* luciferase (RLuc8)(11), a reconstructed ancestral luciferase (Anc^HLD-RLuc^)(13), a fragment-transplanted luciferase (AncFT)(12), and the newly generated luciferase (AncFT-L14) reported herein. (**a**) Multiple sequence alignment with highlighted regions (orange and purple rectangles with black outlines) that were rationally grafted to generate the novel luciferase AncFT-L14. The background color for each residue corresponds to the Clustal X coloring scheme. (**b**) Circular dichroism spectra of all the tested variants. The experiment was performed in 100 mM potassium phosphate buffer (pH 7.5) at 20 °C, and the data are presented as the average of 5 technical replicates. (**c**) Bioluminescence emission spectra of the characterized variants. No spectrum was collected for Anc^HLD-RLuc^ due to the insufficient intensity of its light emission. The experiment was performed in 100 mM potassium phosphate buffer (pH 7.5) at room temperature, and the data are presented as the average of 3 technical replicates. (**d**) Thermal stability analysis to determine the onset temperature (*T*_onset_) and melting temperature (*T*_m_) for all the characterized luciferases. The experiment was performed in 100 mM potassium phosphate buffer (pH 7.5) at a heating rate of 1 °C/min from 20 to 90 °C, and the values are presented as best-fit values ± standard errors. (**e**) Comparison of enzymology parameters, including the catalytic efficiency (*k*_cat_/*K*_m_), product inhibition factor (*K*_p_/*K*_m_), and ‘glow-type effectivity’ (product of the two). The experiment was performed in 100 mM potassium phosphate buffer (pH 7.5) at 37 °C, and the values are presented as best-fit values ± standard errors. (**f**) Bioluminescence decay measurements in cell lysates after overexpression of the characterized variants. No decay curve was collected for Anc^HLD-RLuc^ due to insufficient light emission intensity. The experiment was performed in 100 mM potassium phosphate buffer (pH 7.5) at 37 °C, and the data are presented as the average of 3 independent replicates.

Most importantly, enzymological and biophysical analyses revealed substantial improvements in both catalytic parameters and light-emission properties for the engineered variant. AncFT-L14 displayed 1.3-, 2.4-, and 16,300-fold higher catalytic efficiency (*k*_cat_/*K*_m_) than those of RLuc8, AncFT, and the original ancestral enzyme Anc^HLD-RLuc^, respectively. This was accompanied by a pronounced 15.9-, 1.6-, and 45.0-fold increase in the product inhibition factor (*K*_p_/*K*_m_), respectively (**Figure 5e, Figure S4**). This shift indicates the AncFT-L14 enhanced substrate preference over the product (increased *K*_p_/*K*_m_ ratio), a result consistent with the potential targeted acceleration of the rate-limiting, product-release-associated conformational step. To evaluate luciferase performance in biotechnologically relevant luminescence assays, we combined both parameters into a composite metric *k*_cat_/*K*_m_ × *K*_p_/*K*_m_, herein termed ‘glow-type effectivity’. This metric directly reflects the ability to maintain sustained light output under turnover conditions despite product accumulation. By this measure, AncFT-L14 outperformed AncFT nearly 4-fold, RLuc8 by more than 20-fold, and Anc^HLD-RLuc^ by more than 700,000-fold (**Figure 5e**).

In accordance with its superior catalytic profile, AncFT-L14 exhibited prolonged luminescence and enhanced emission stability in cell lysates, surpassing not only RLuc8 but also AncFT, a previously established glow-type bioluminescence benchmark (**Figure 5f**). Together, these results demonstrate that mechanism-guided rational loop grafting can successfully bypass kinetic bottlenecks to yield a luciferase with superior catalytic efficiency and stable glow-type bioluminescence. These properties make AncFT-L14 an excellent candidate for applications that require prolonged or continuous acquisition of the luminescence signal.

### Integrative analysis corroborates altered conformational dynamics as the molecular basis of the enhanced bioluminescence

To uncover the molecular origins underlying the improved performance and enhanced glow-type bioluminescence of AncFT-L14, we performed an integrative analysis combining transient kinetics (**Figure 6**), structural biology (**Figure 7**), and *in silico* modeling (**Figure 8**). This multi-modal approach was designed to distinguish whether the engineered improvements arise from changes in chemical catalysis, ligand binding and positioning, or fundamental changes in protein conformational dynamics.

**Figure 6:**
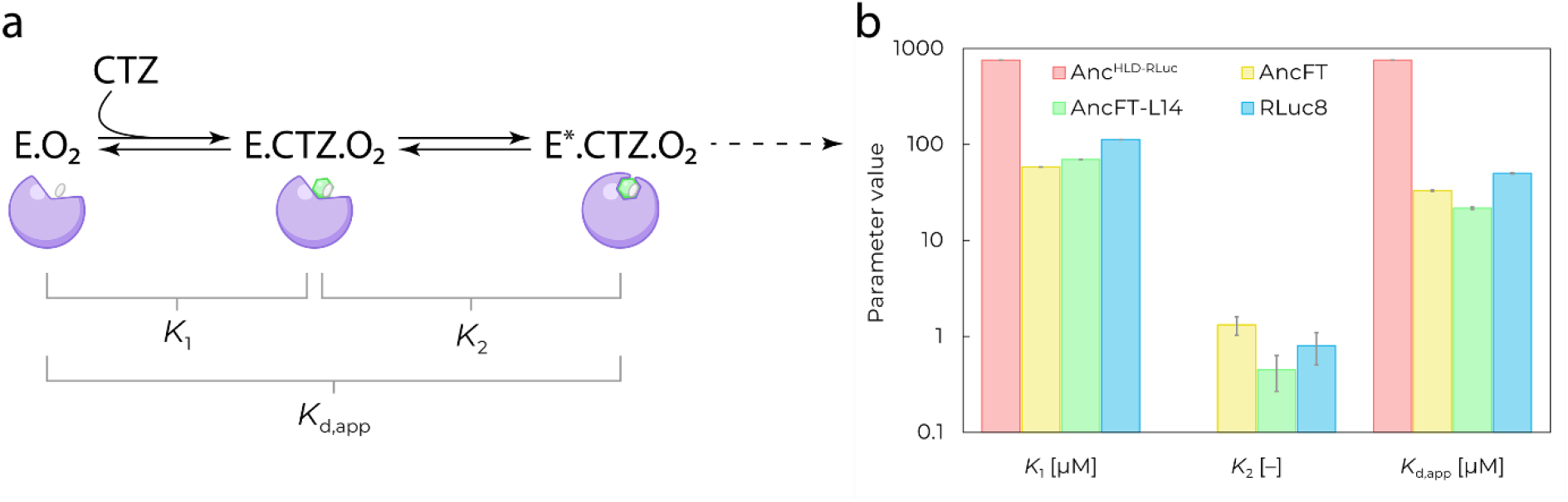
Transient kinetic analysis of the induced-fit transport process steps for the characterized variants Anc^HLD-RLuc^, AncFT, AncFT-L14, and RLuc8. (**a**) Illustrative representation of the two steps involved in the induced-fit binding mechanism and the influence of the corresponding equilibrium constants on each of these steps. E.O_2_ = enzyme–oxygen complex; CTZ = coelenterazine substrate; E.CTZ.O_2_ and E^*^.CTZ.O_2_ = ternary enzyme–coelenterazine–oxygen complex in two different conformations. The nomenclature and illustrations correspond to the symbols used in **Figure 3**. (**b**) Comparison of equilibrium constants characterizing individual steps of the induced-fit binding mechanism. The *K*_2_ value is not provided for Anc^HLD-RLuc^ because of its rigid, simple one-step binding and no conformational change observed upon initial collision. Note the logarithmic scale of the y-axis. The experiment was performed in 100 mM potassium phosphate buffer (pH 7.5) at various temperatures (9 to 22 °C), and the values are presented as best-fit values ± standard errors.

**Figure 7:**
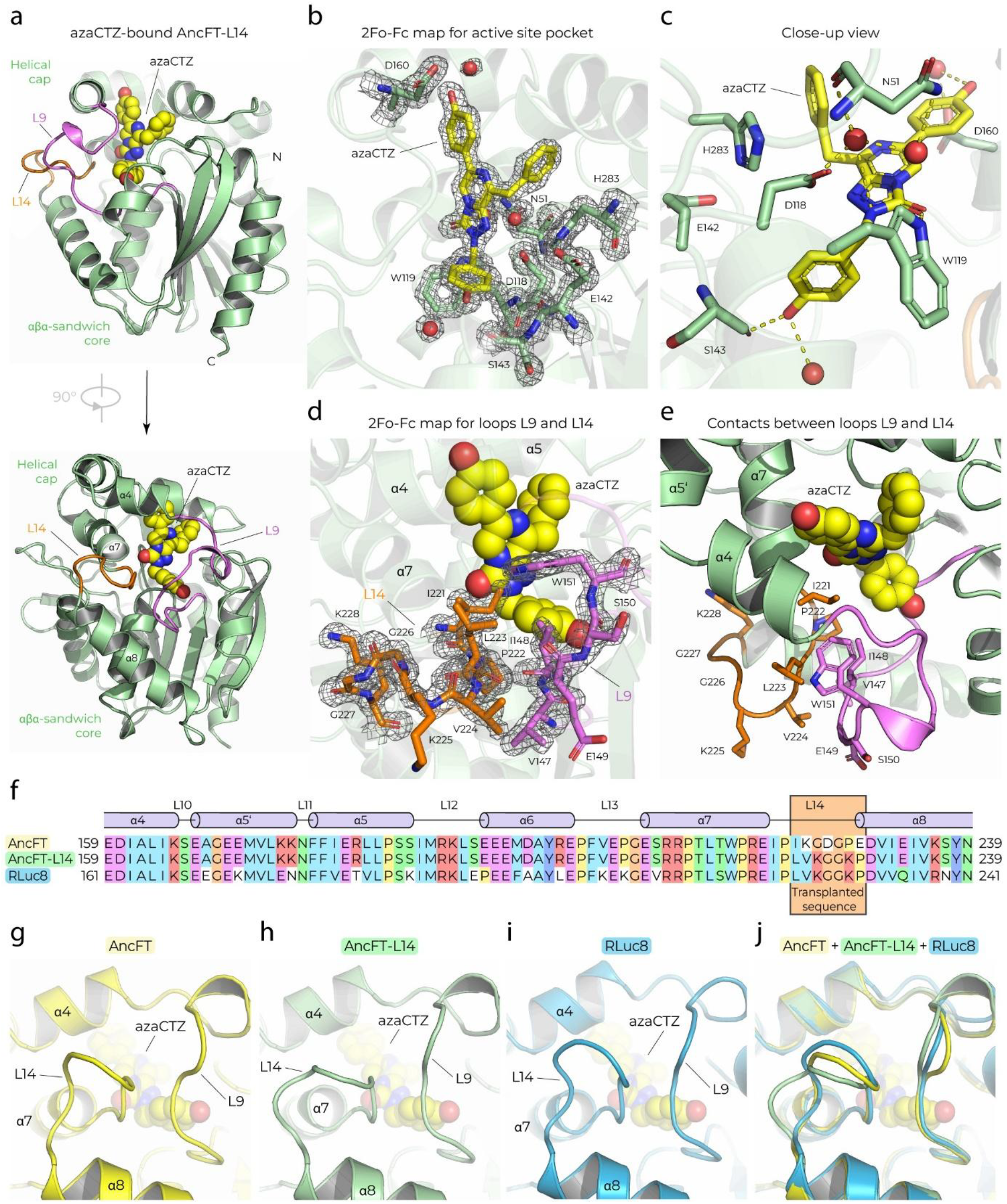
Co-crystal structure of the luciferase AncFT-L14 in complex with azacoelenterazine (azaCTZ). (**a**) Cartoon representation of the overall structure of the AncFT-L14 (PDB ID: 7OME) enzyme (green) with bound azaCTZ (spheres, with carbon atoms in yellow). The loop L9 is colored violet, and the loop L14 is colored orange. (**b**) 2Fo-Fc electron density (contour level 1.5 σ) for azaCTZ (yellow sticks) and selected residues (green sticks) at the active site pocket. (**c**) Close-up view of azaCTZ bound in the AncFT-L14 active site. Residues creating the active site are shown in green stick representation, azaCTZ is shown in yellow stick representation, and key hydrogen bonds are shown as dashed yellow lines. (**d**) 2Fo-Fc electron density (contour level 1.5 σ) for the loop L14 (orange sticks) and a partial sequence (V_147_IESW) of the loop L9 (violet sticks). AzaCTZ is shown as yellow spheres. (**e**) Close-up view of the molecular contacts between the loops L9 (violet) and L14 (orange). (**f**) Structure-based partial sequence alignment of AncFT, AncFT-L14, and RLuc8. The secondary structure elements are shown above the alignment. The L14 sequence of RLuc8 used to replace that of AncFT, resulting in AncFT-L14, is labeled with an orange box. (**g**-**j**) Structural comparisons of the loops L9 and L14 conformations in AncFT (PDB ID: 7QXR) (g), AncFT-L14 (PDB ID: 7OME) (h), and RLuc8 (PDB ID: 2PSF) (i), and superposition of these loops (j). Note that the loop L14 in AncFT-L14 adopts a unique conformation.

**Figure 8:**
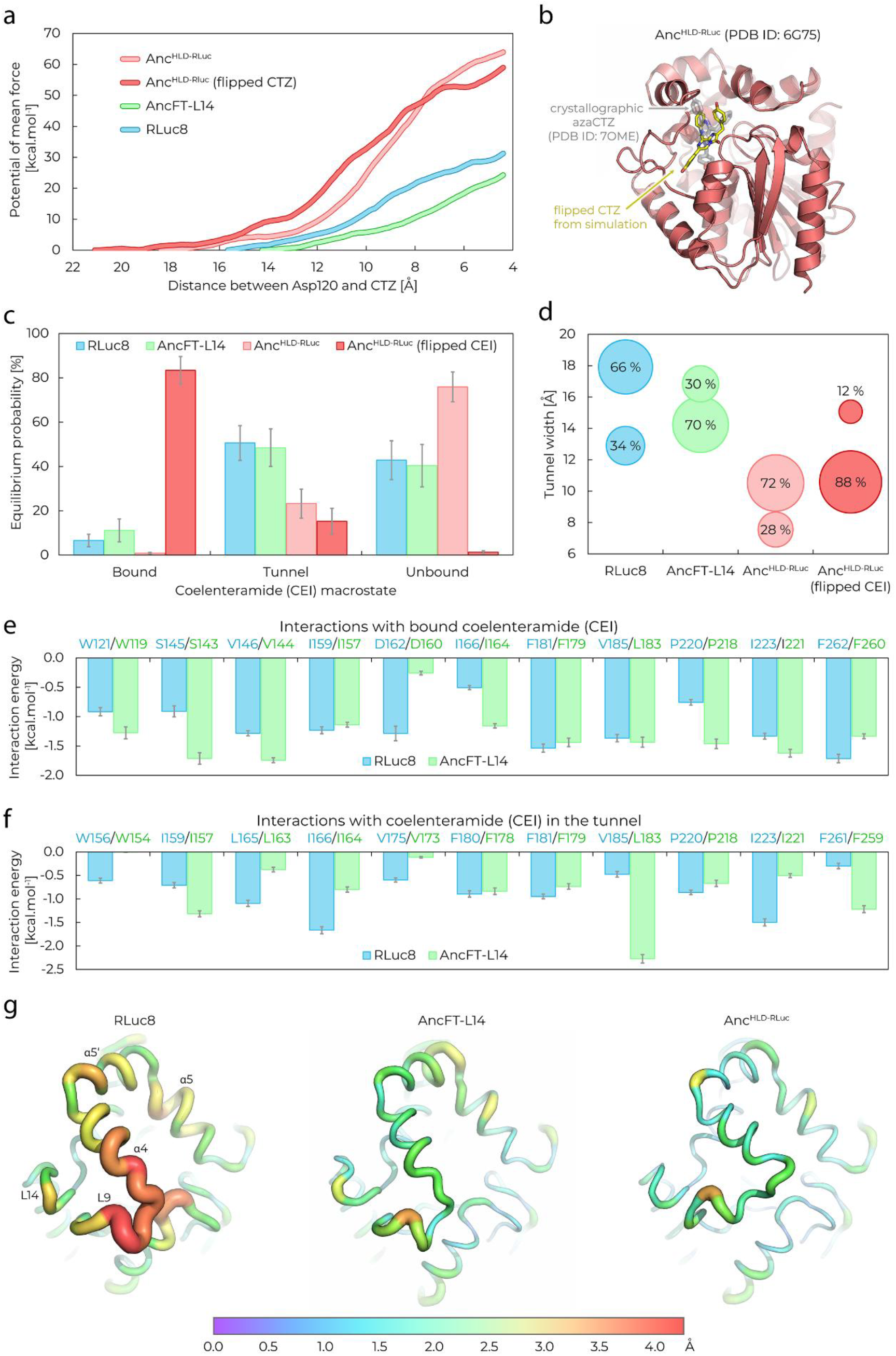
Computational simulations of enzyme–ligand complexes of RLuc8, AncFT-L14, and Anc^HLD-RLuc^. (**a**) Potential of mean force used to pull coelenterazine (CTZ) into the active site of the luciferases obtained by adaptive steered molecular dynamics (ASMD). (**b**) A flipped CTZ end pose (yellow sticks) from simulated binding to Anc^HLD-RLuc^ (red cartoon). The crystallographic ligand obtained for AncFT-L14 is shown as a reference (translucent gray sticks, PDB ID: 7OME). (**c**) Equilibrium probabilities of coelenteramide (CEI) macrostates constructed on the basis of the CEI-active site distance (‘bound’, ‘tunnel’, ‘unbound’) derived from the adaptive sampling of CEI unbinding. The simulations started from the crystal-like CEI pose for each enzyme and additionally, from the flipped pose shown in (b) in Anc^HLD-RLuc^. The error bars indicate the standard error of the equilibrium probability obtained by bootstrapping a random 80 % of the data 100 times. (**d**) Analysis of access tunnel opening potential during the adaptive sampling of CEI unbinding. The macrostates (shown as bubbles) were constructed based on the width of the tunnel mouth; the equilibrium probabilities of these macrostates are written as percentages inside the corresponding bubbles and reflected in the bubble size. (**e**–**f**) The interaction energies of CEI with each residue of the ‘bound’ CEI macrostate (e) and the ‘tunnel’ macrostate (f) from adaptive sampling with RLuc8 and AncFT-L14. For clarity, only attractive interactions are shown (less than −1.0 kcal.mol^−1^ for the ‘bound’ states and less than −0.5 kcal.mol^−1^ for the ‘tunnel’ states). The error bars indicate the standard errors. Mapping of the interacting residues on the corresponding structures is provided in **Figures S18 and S19**. (**g**) Cap domains of RLuc8, AncFT-L14, and Anc^HLD-RLuc^ with mapped root-mean-square fluctuations (RMSFs) of Cα atoms in the ‘unbound’ states from adaptive sampling.

### Transient kinetic analysis revealed that loop grafting selectively accelerated induced-fit dynamics

Pre-steady-state kinetic characterization enabled us to dissect the observed improvement in AncFT-L14 catalytic efficiency into individual kinetic steps (**Figures S5** and **S6, Table SIII, Supplementary Note 2**). The results showed that enhancement originates almost exclusively from modulation of protein conformational flexibility (**Figure 6**). While the initial enzyme–ligand collision step was only marginally affected (*K*_1_ shows a 1.3-fold difference compared to AncFT), the subsequent conformational transition displayed a substantially more favorable equilibrium, indicating selective stabilization of the catalytically competent closed state (*K*_2_ shows a 3.0-fold difference compared to AncFT). These findings confirm that mechanism-based engineering precisely modified the targeted induced-fit conformational rearrangement without substantially altering other steps. Given that rational engineering of the induced-fit/conformational step is a challenging task rarely reported in the literature(*28–32*), this result underscores the power of the loop-grafting strategy and the LoopGrafter tool(*26*) to rationally design biocatalysts with tailored kinetic properties.

### Co-crystal structure uncovered preserved substrate binding but unique loop conformational organization

To assess whether loop grafting alters active-site architecture, we solved the co-crystal structure of AncFT-L14 in complex with the non-oxidizable substrate analog azacoelenterazine (azaCTZ)(*14, 33*) at 1.5 Å resolution (**Figure 7, Table SIV, Supplementary Note 3**). The overall azaCTZ-binding mode and its key molecular contacts with active-site residues remained very similar, if not identical, to those of the previously characterized AncFT–azaCTZ luciferase complex (**Figure 7a–c, Figure S7**)(*14*), suggesting that substrate positioning and chemical activation remained unchanged. In contrast, the grafted loop L14 adopted an atypical conformation in AncFT-L14, distinguishing it from AncFT and RLuc8 (**Figure 7d–j**). Specifically, the transplanted segment L_223_VKGGKP is positioned near the bound substrate analog (azaCTZ), but does not form extensive direct physical contacts with it. Instead, the loop L14 makes numerous, mostly nonpolar and hydrophobic contacts with the neighboring loop L9. These structural features of enhanced inter-loop complementarity and L9–L14 interface plasticity suggest that loop grafting reshapes the conformational landscape surrounding the access tunnel rather than the chemical machinery of the active site. Such coordinated loop motions appear to be crucial for facilitating substrate loading and product unloading during catalysis via the enzyme conformational changes identified in our kinetic analysis.

### Refined structural analysis of the active site identified further molecular details of substrate and oxygen positioning

By determining the structure at a higher resolution (1.5 Å) than previously reported structures of *Renilla*-type luciferases, we unambiguously resolved two water molecules at the bottom of the active-site pocket, proximal to the reactive C2 carbon atom of the native CTZ (**Figure 7b-c**). This position is consistent with the putative dioxygen-binding site and mimics oxygen coordination within the Michaelis complex (**Supplementary Note 3**). The observed geometry highlights the potential contributions of D118, N51, and W119 in properly orienting the dioxygen co-substrate for nucleophilic attack by the C2 carbon of the activated CTZ. In addition, the side chain of W119 forms hydrogen bonding with the carbonyl group of azaCTZ, providing an additional anchoring point to stabilize the enzyme–substrate complex (**Figure 7c, Figure S8**). Importantly, these structural features are conserved relative to previously characterized *Renilla* luciferases, indicating that the enhanced performance of AncFT-L14 does not arise from alterations in substrate positioning or transient state formation. Instead, the improved catalytic profile is a direct result of the optimized geometry and coordinated conformational flexibility of loops, including the transplanted loop L14.

### In silico modeling linked loop reorganization to altered ligand exchange dynamics and access tunnel closing/opening

To further evaluate whether loop grafting affected protein tunnel dynamics and ligand transport efficiency, we performed additional molecular docking (**Figures S9**– **S11, Table SV**) and molecular dynamics (**Figure 8**) simulations. Specifically, we analyzed CTZ substrate binding using adaptive steered molecular dynamics (ASMD; **Supplementary Note 4**), and CEI product release using adaptive sampling molecular dynamics (**Supplementary Notes 5**). Consistent with our kinetic and structural observations, substrate binding to AncFT-L14 required 7 kcal.mol^−1^ less work than binding to RLuc8 (**Figure 8a, Figures S12** and **S13**). Product release simulations (**Figures S14–S17**) also revealed that AncFT-L14 exhibited stronger interactions with CEI compared to RLuc8 (**Figure 8c, Table SVI** and **SVII**). Importantly, AncFT-L14 maintained strong ligand interactions mostly within the catalytic active site (**Figure 8e, Figure S18**) and favored the catalytically competent conformation with the closed tunnel (**Figure 8d**) which effectively avoids the nonproductive trapping of ligands within the access tunnel. In striking contrast, RLuc8 exhibited stronger ligand interactions predominantly with the access tunnel-lining residues located in the dynamic loop L9 and α4 fragment (**Figure 8f Figure S19**). This dynamic element was moving away to the side during CEI release (**Figure 8g**), overall resulting in a wide, open-tunnel conformation in which RLuc8 remained for most of the snapshots (**Figure 8d**). These changes in ligand trafficking dynamics and tunnel closing/opening provide a molecular rationale for the enhanced turnover efficiency and prolonged emission stability observed experimentally in AncFT-L14.

### Simulations with the Anc^HLD-RLuc^ provided additional insight into the relationship between ligand orientation, product inhibition, and emission behavior

Apart from a deeper exploration of AncFT-L14, simulations of the original ancestral protein, Anc^HLD-RLuc^, also yielded unexpected, previously unseen insights when compared to all other luciferases (**Supplementary Notes 4** and **5**). ASMD revealed that CTZ substrate binding in Anc^HLD-RLuc^ preferentially adopted a flipped, nonproductive orientation (**Figure 8b**) requiring twice the energy expenditure compared to binding in AncFT-L14 or RLuc8 (**Figure 8a**). Similarly, simulations of CEI product release—in the orientation captured with RLuc8—yielded a considerably lower affinity for Anc^HLD-RLuc^, as demonstrated by the 90-fold lower probability of the ‘bound’ state than the ‘unbound’ state (**Figure 8c, Table SVI**). Strikingly, simulations with the flipped CEI showed a pronounced stabilization of the Anc^HLD-RLuc^–CEI complex by yielding a markedly higher probability of the ‘bound’ state relative to any other simulated protein– ligand pair (**Figure 8c, Table SVI** and **SVII**). This tight, nonproductive mode of binding explains the strong product inhibition reported for Anc^HLD-RLuc^ previously(*12*) and contrasts sharply with the AncFT-L14 preference for productive ligand orientations. Since product inhibition has been linked to ‘flash-type’ bioluminescence, the absence of this flipped inhibitory binding in AncFT-L14 provides additional molecular evidence of its ‘glow-type’ bioluminescence. This outcome further implies that the control of ligand orientation and conformational gating, rather than substrate chemistry alone, is the critical determinant of luciferase performance.

Altogether, the convergent kinetic, structural, and computational analyses demonstrate that the improved catalytic and bioluminescence performance of AncFT-L14 arises from the selective reshaping of protein conformational dynamics that govern ligand exchange, without perturbing substrate chemical conversion or introducing unfavorable trade-offs. This integrative mechanistic framework not only explains the emergence of glow-type behavior but also validates loop grafting as an effective strategy for engineering enzyme dynamics to achieve significant functional gains.

## Discussion

In addition to being remarkable light-emitting machines in nature, luciferases have become invaluable tools in research and clinical diagnostics as bioluminescence reporters and bioimaging agents(*4–6*). Despite their enduring popularity, the molecular principles governing luciferase efficiency and light emission features have only recently begun to unfold but remain incompletely understood. Here, we provide the comprehensive mechanistic determinants of *Renilla* luciferase (RLuc) catalysis by integrating steady-state and transient kinetics with structural biology and *in silico* simulations. This analysis reveals that RLuc performance is constrained by multiple interdependent factors, including oxygen undersaturation, oxygen-induced irreversible inactivation, and a rate-limiting induced-fit conformational transition during product release. We further demonstrate how selectively targeting protein dynamics, rather than chemistry, can overcome these intrinsic limitations.

Our extended steady-state kinetic analysis fundamentally revises the long-standing view of *Renilla* luciferase efficiency under physiological conditions. We show that the enzyme operates far below oxygen saturation due to a high Michaelis constant for the oxygen co-substrate, leading to a significant and systematic underestimation of the true turnover number *k*_cat_ in all previous studies(*11, 12, 14*). In light of that, while increasing oxygen affinity might appear to be a straightforward engineering solution, our results at high concentrations of dissolved oxygen expose an intrinsic trade-off: elevated oxygen levels accelerate irreversible enzyme inactivation, likely mediated by reactive radical intermediates inherent to the catalytic mechanism(*14*). These findings suggest that luciferase evolution and function are shaped by a delicate balance between catalytic activity and oxidative self-damage, imposing limitations that cannot be alleviated by engineering oxygen binding alone. Similar constraints may be broadly relevant among oxygen-activating enzymes that rely on radical intermediates, including cytochrome P450s, flavin-dependent monooxygenases, and non-heme iron oxygenases(*34–38*).

Interestingly, similar irreversible enzyme inactivation after a given number of turnover cycles has been reported for native RLuc-wt(*20*), NanoKAZ/NanoLuc(*21*), and Gaussia luciferase (GLuc)(*19*), but never for RLuc8 under physiological conditions, suggesting that the consensus-based mutagenesis, that improves RLuc8 resistance(*11*), also decreased its susceptibility to inactivation. Pronounced inactivation with increasing oxygen concentration clearly points toward an oxygen-induced mechanism of inactivation, which is in accordance with the previously reported detection of 17 Da adducts for inactivated GLuc(*22*). These observations are in line with the recently deciphered catalytic mechanism of CTZ bioluminescence, which proceeds via highly reactive oxygen radical intermediates(*14*) that are likely the source of enzyme covalent modification and inactivation. While compatible with luciferase roles in nature, this inactivation adversely affects bioimaging applications with extended acquisition times.

In addition to oxygen-associated limitations, our transient kinetic analysis identifies the major bottleneck in the overall catalytic cycle of RLuc8: the conformational opening of the product-bound enzyme, rather than substrate binding or chemical transformation, is the major rate-limiting step under the assay conditions. This insight reframes luciferase catalysis as a process limited primarily by protein dynamics. Importantly, it also identifies a new mechanism-grounded engineering target that avoids the inherent liabilities associated with modifying oxygen affinity. However, selectively targeting conformational transitions without impairing the chemical transformation or the initial enzyme–ligand collision remains a formidable challenge with very few documented success cases to date(*28–32*). We addressed this issue by employing rational loop grafting to reshape the conformational energy landscape of the enzyme. By exploiting this principle, we demonstrate that directed modulation of induced-fit dynamics can yield substantial improvements in both luciferase catalytic efficiency and light emission stability.

Detailed integrative analysis of the resulting improved variant AncFT-L14 revealed that engineering of dynamic loops surrounding the active site selectively accelerated the targeted conformational flexibility. Kinetic characterization demonstrated that the conformational transition of the ligand-bound enzyme was enhanced almost exclusively, while the initial ligand collision step of induced-fit binding remained nearly unaffected. Structural analysis confirmed the preservation of the active-site architecture, while discovering novel inter-loop interactions that alter access-tunnel organization and substrate/product (un)loading. Molecular dynamics simulations further showed that these changes redirect ligand trafficking pathways, favor productive binding and release, and reduce nonproductive trapping, thus providing a direct physical basis for the observed glow-type bioluminescence. Beyond the engineered variant AncFT-L14, simulations with the ancestral luciferase Anc^HLD-RLuc^ uncovered an unexpected strong preference for a nonproductive, flipped ligand orientation whose expulsion from the active site required a substantial amount of energy. This energetic barrier provides a molecular rationale for the pronounced product inhibition reported for Anc^HLD-RLuc^ previously(*12*). All these new insights present a mechanistic foundation for further design of luciferases with emission profiles tailored to specific applications.

## Conclusions

In conclusion, by resolving the kinetic bottlenecks identified in this work, our study establishes a mechanism-guided pathway for engineering *Renilla*-type luciferases with enhanced catalytic efficiency and glow-type bioluminescence. This was demonstrated by generating the loop-grafted variant AncFT-L14, which exhibits a substantially more stable light emission signal, surpassing the *Renilla*-type enzymes used in current state-of-the-art bioluminescence assays. By integrating enzyme kinetics, structural biology, and molecular dynamics, we provide a blueprint for rationally designing enzymes in which conformational control, rather than chemical reactivity or ligand positioning, is the primary determinant of performance. Our work therefore underscores protein conformational dynamics as a tunable and underexplored axis of enzyme engineering. While most engineering strategies focus on optimizing active-site chemistry or substrate affinity(*39, 40*), our results demonstrate that selectively reshaping induced-fit processes can deliver substantial functional gains while avoiding unfavorable side effects. This concept is likely applicable beyond luciferases, particularly for enzymes whose catalytic cycles involve buried active sites and gated ligand exchange(*41–44*). We anticipate that these insights will guide the development of next-generation bioluminescent tools and navigate broader efforts to engineer enzyme dynamics for functional innovation.

## Data availability

Atomic coordinates and structure factors have been submitted to the Protein Data Bank (www.wwpdb.org)(*45*) under the PDB ID accession code 7OME. Molecular dynamic simulation data were deposited in the Zenodo repository with the assigned DOI of 10.5281/zenodo.15516240.

## Acknowledgements

This research was supported by the European Union’s Horizon 2020 Research and Innovation Programme under grant agreement Nos. 857560 and 101136607, and by the project National Institute for Neurology Research (No. LX22NPO5107 MEYS), financed by the European Union – Next Generation EU. The authors thank the RECETOX Research Infrastructure (No. LM2023069) financed by the Ministry of Education, Youth, and Sports. Computational resources were provided by the e-INFRA CZ and ELIXIR-CZ (Nos. 90254 and LM2023055), supported by the Ministry of Education, Youth and Sports of the Czech Republic. The work reported in this paper was supported by the Czech Science Foundation (GX25-17329X). Martin Toul was supported by the Brno Ph.D. Talent scholarship (JCMM & Brno City Municipality) and the Fulbright Postgraduate Scholarship (Fulbright Commission). Research in Pimchai Chaiyen laboratory is supported by the NSRF via the Research and Innovation Acceleration Agency for Competitiveness and Area Development (RCAD) – Program Management Unit for Technology and Innovation for Future Industries (PMU-B): Brainpower for Future Industries (No. B38G690046). Crystallographic data were collected at the PXIII beamline at the Swiss Light Source (SLS) in Villigen (Switzerland). We are grateful to the members of the SLS synchrotron for the use of their beamline and help during data collection. We acknowledge CF Biomolecular Interactions and Crystallography of CIISB, Instruct-CZ Centre, supported by MEYS CR (LM2023042) and European Regional Development Fund-Project „Innovation of Czech Infrastructure for Integrative Structural Biology” (No. CZ.02.01.01/00/23_015/0008175).

