## Supporting information for "Conformational gating of product release sets the catalytic ceiling of a bioluminescent reporter"

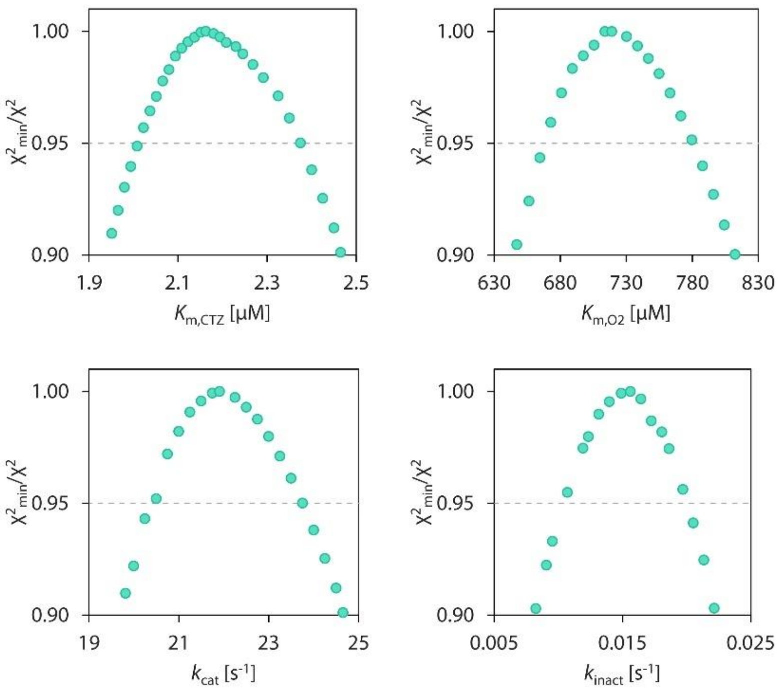

**Supplementary Fig. 1. Confidence contour analysis of parameters obtained during the RLuc8 oxygen-dependent steady-state kinetic data analysis.** All the parameters are well constrained by the collected kinetic data, confirming the robustness of the performed analysis. The grey dashed lines represent the $x^2$ threshold of 0.95. The notation of parameters corresponds to those illustrated in Fig. 2 in the main text.

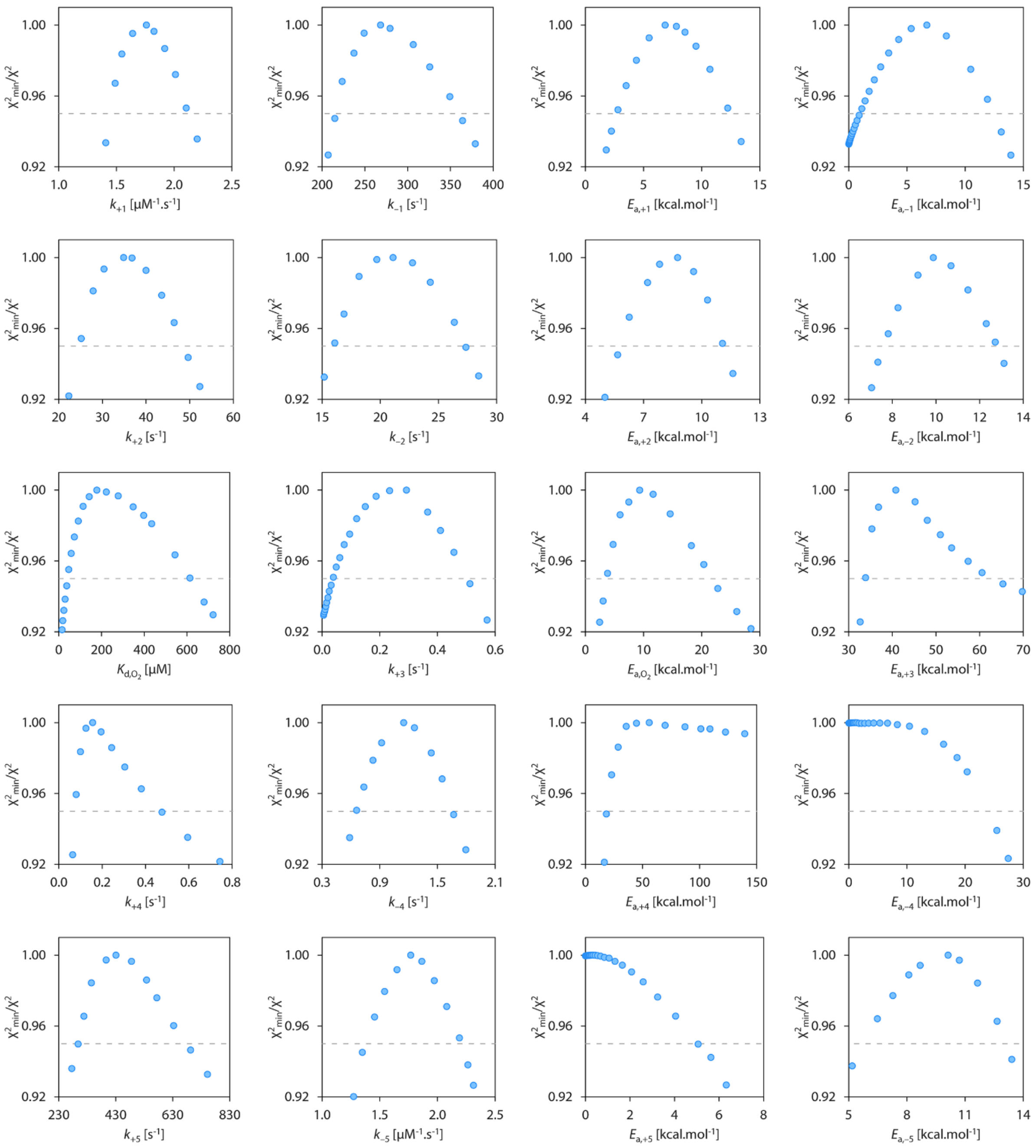

**Supplementary Fig. 2. Confidence contour analysis of parameters obtained during the global analysis of RLuc8 pre-steady-state transient kinetic data.** All the parameters are well constrained by the collected kinetic data, confirming the robustness of the performed analysis. The grey dashed lines represent the $x^2$ threshold of 0.95. The notation of parameters corresponds to those illustrated in Fig. 4 in the main text.

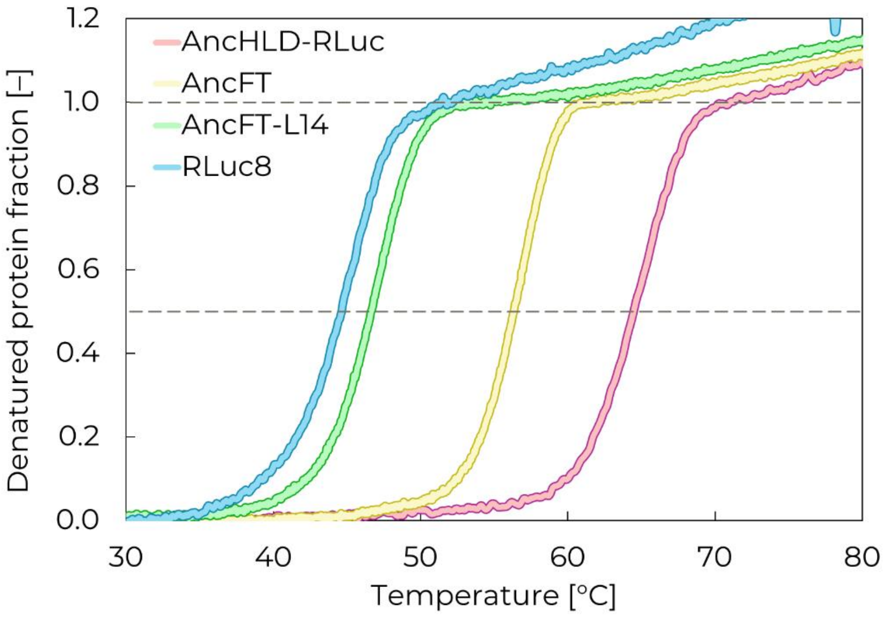

**Supplementary Fig. 3. Melting curves monitoring thermal denaturation of characterized luciferases RLuc8, Anc$^{HLD\text{-}RLuc}$, AncFT, and AncFT-L14.** Dashed lines indicate the state of half-denatured (corresponding to the $T_m$ value) and fully-denatured proteins, respectively. The experiment was performed in 100 mM potassium phosphate buffer pH 7.5 and the results are shown as an average of three technical replicates.

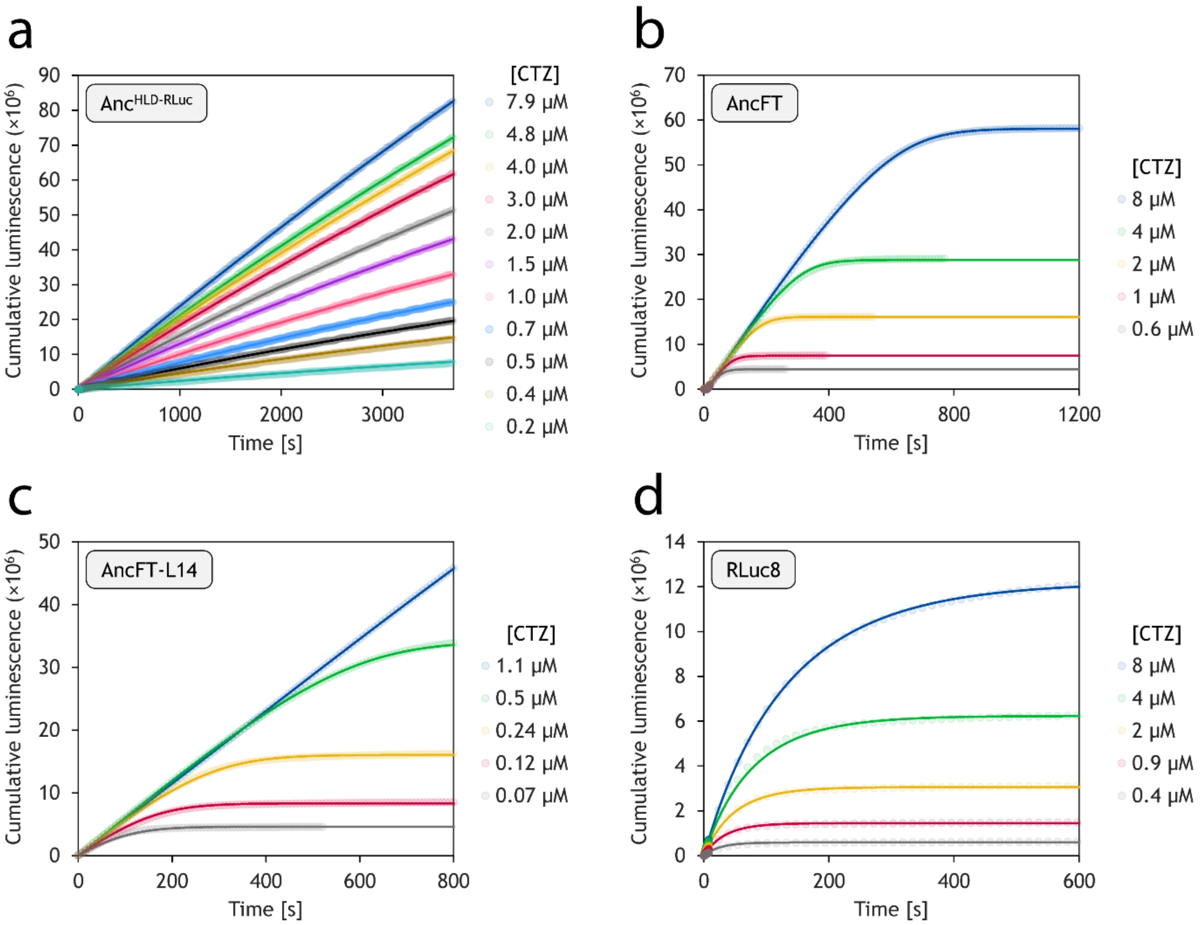

**Supplementary Fig. 4. Numerical integration analysis of steady-state kinetic data collected for characterized luciferase variants Anc$^{HLD-RLuc}$ (a), AncFT (b), AncFT-L14 (c), and RLuc8 (d).** Dotted lines correspond to experimental data while solid lines represent the best fit according to the state-state model provided in the Online Methods section. The experiment was performed in 100 mM phosphate buffer pH 7.5 at 37 °C and the data are shown as an average of three technical replicates. The resulting concentrations after mixing were 640 nM Anc$^{HLD-RLuc}$, 130 nM AncFT, 6 nM AncFT-L14, and 17 nM RLuc8. The concentrations of the coelenterazine substrate varied from 0.07 to 8 µM and are stated in the figure legend.

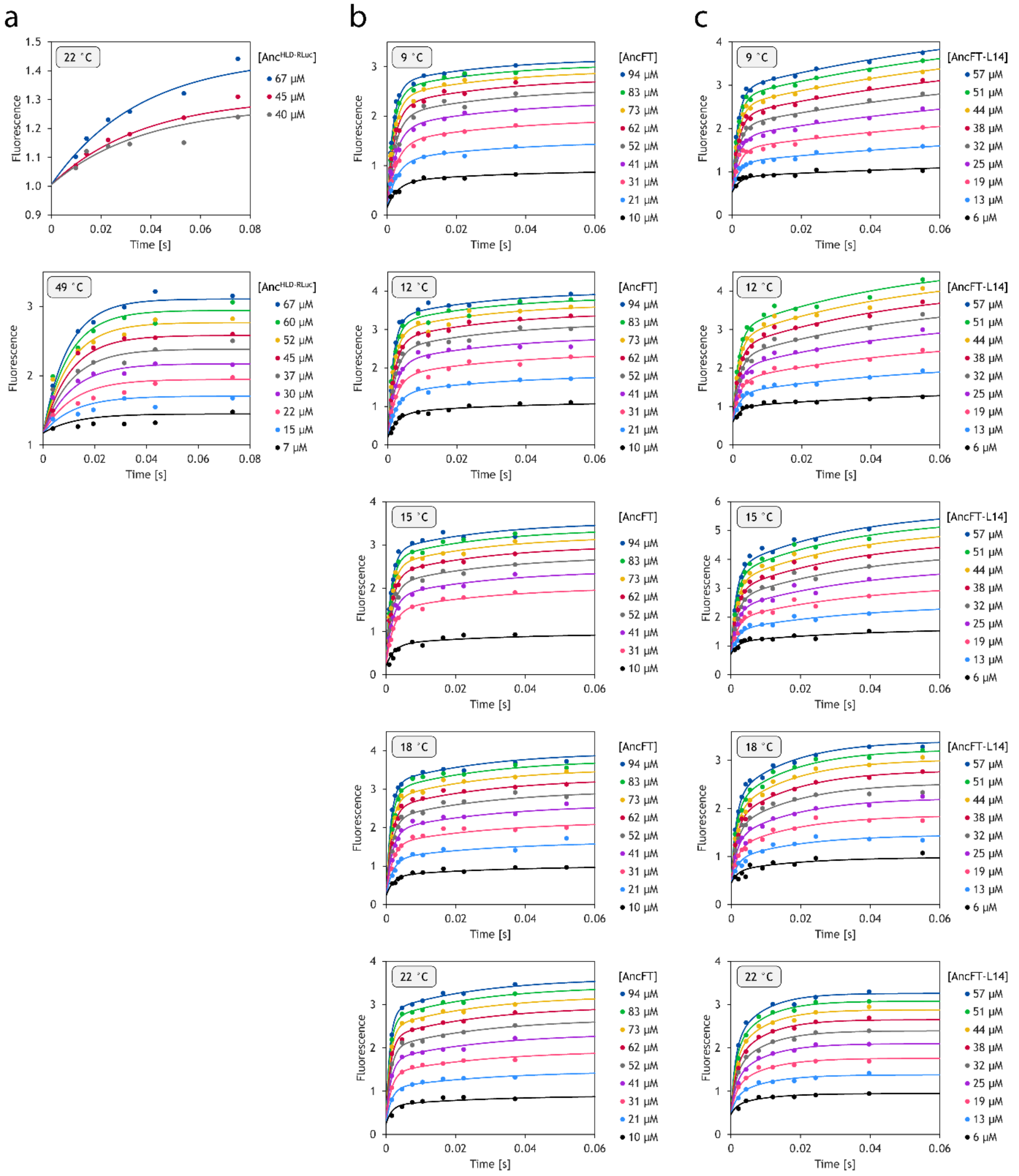

**Supplementary Fig. 5. Numerical integration analysis of on-chip transient kinetic fluorescence data collected for the tested variants Anc$^{HLD\text{-}RLuc}$ (a), AncFT (b), and AncFT-L14 (c).** Dots correspond to experimental data while solid lines represent the best fit according to the induced-fit model provided in Fig. 6a in the main text. The experiment was performed in 100 mM phosphate buffer pH 7.5 at varying temperatures stated in the top-left corner of each graph. The resulting concentrations after mixing were 7-67 µM Anc$^{HLD\text{-}RLuc}$, 10-94 µM AncFT, and 6-57 µM AncFT-L14 while the coelenterazine concentration was fixed at 30 µM.

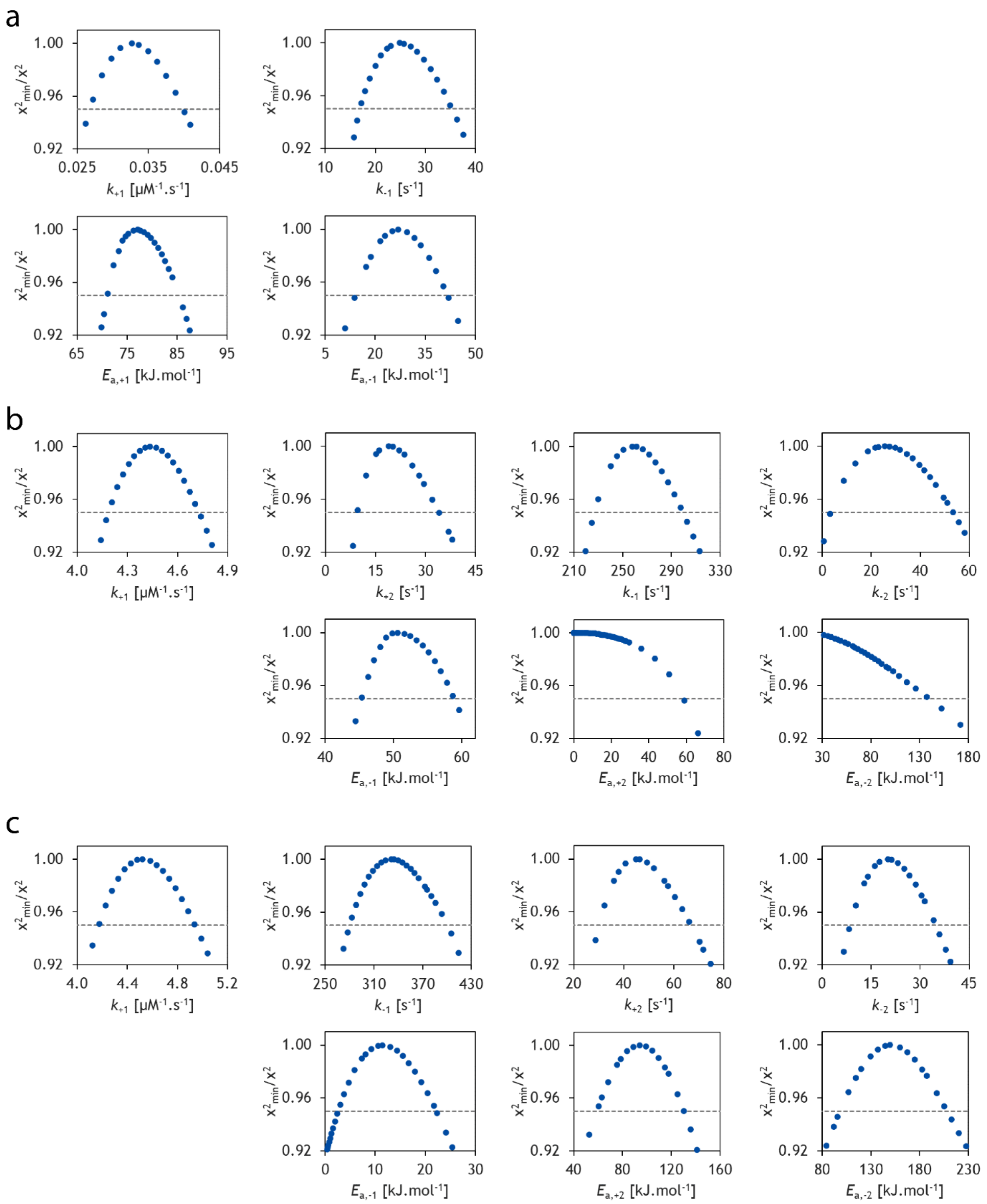

**Supplementary Fig. 6. Confidence contour analysis of rate constants and activation energies of the induced-fit step determined for the analyzed variants Anc$^{HLD\text{-}RLuc}$ (a), AncFT (b), and AncFT-L14 (c).** The grey dashed lines represent the $x^2$ threshold of 0.95. The notation of parameters corresponds to those presented in Supplementary Table III. Confidence contour analysis was based on the data fitting shown in Supplementary Fig. 5. Data interpretation is provided in Supplementary Note 2.

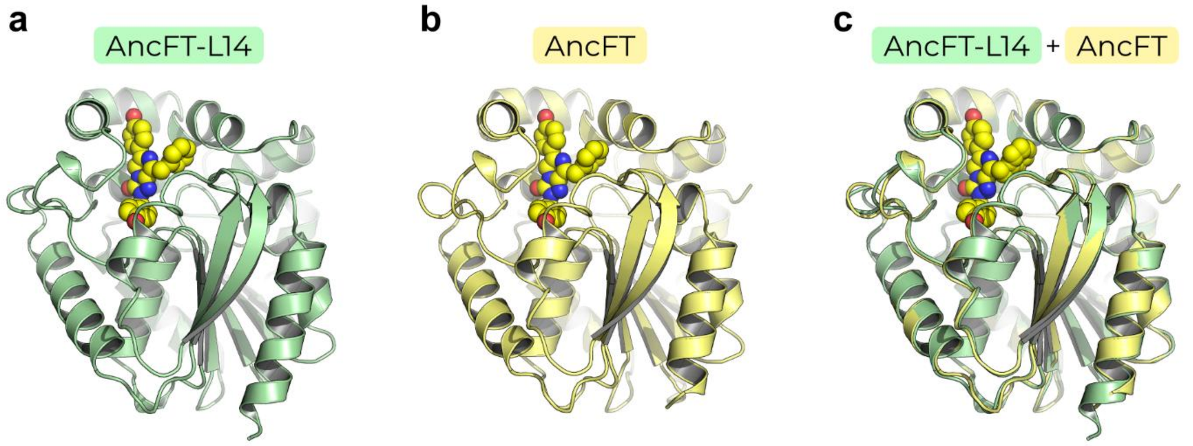

**Supplementary Fig. 7. Structural comparison of AncFT-L14 (PDB ID: 7OME) and AncFT (PDB ID: 7QXR).** Cartoon representation of azacoelenterazine (azaCTZ)-bound AncFT-L14 (a) and AncFT (b), and their superposition (c). The azaCTZ molecule is shown as a yellow space-filling calotte model.

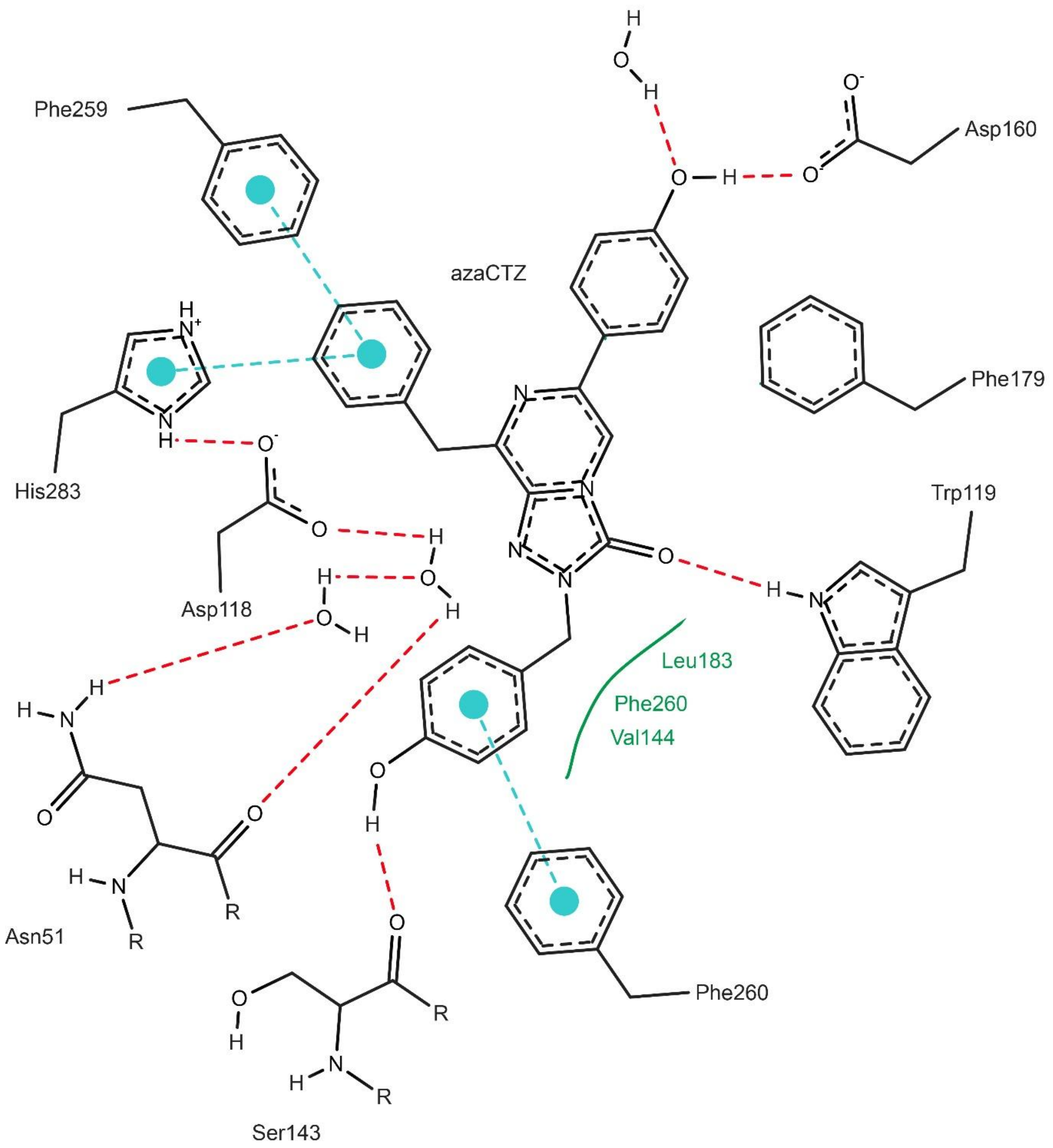

**Supplementary Fig. 8. Two-dimensional representation of molecular contacts between AncFT-L14 amino acid residues and water molecules interacting with azacoelenterazine (azaCTZ).** Hydrogen bonds are depicted as red dotted lines, π-π interactions as blue dotted lines. The figure was generated with PoseView(1).

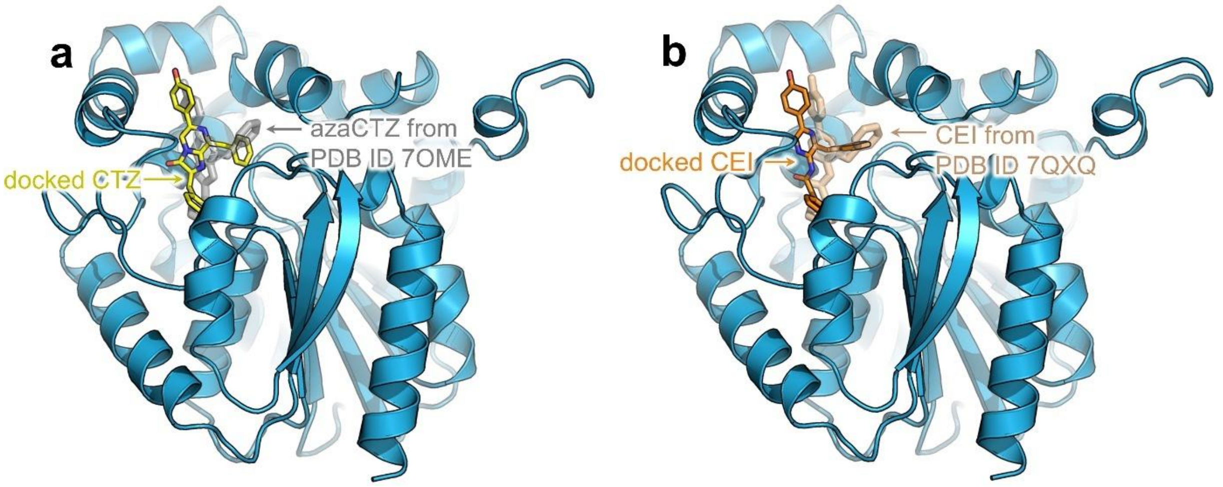

**Supplementary Fig. 9. Molecular docking of coelenterazine (CTZ) and coelenteramide (CEI) to the structure of RLuc8 (PDB ID: 2PSF) shown as a blue cartoon.** (**a**) The predicted pose of CTZ shown as yellow sticks, and the crystallographic pose of azacoelenterazine (azaCTZ) as transparent grey sticks. (**b**) The predicted pose of CEI shown as orange sticks, and the crystallographic pose as transparent brown sticks. Structure 7OME depicts the azaCTZ-bound AncFT-L14 luciferase and 7QXQ the CEI-bound AncFT luciferase.

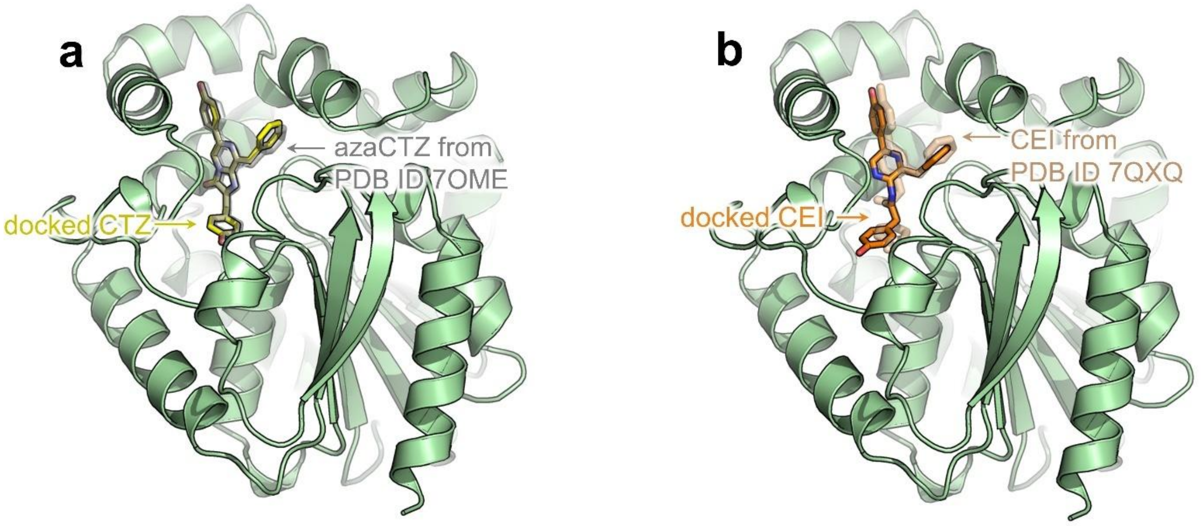

**Supplementary Fig. 10. Molecular docking of coelenterazine (CTZ) and coelenteramide (CEI) to the structure of AncFT-L14 (PDB ID: 7OME) shown as a light green cartoon.** (**a**) The predicted pose of CTZ shown as yellow sticks, and the crystallographic pose of azacoelenterazine (azaCTZ) as transparent grey sticks. (**b**) The predicted pose of CEI shown as orange sticks, and the crystallographic pose as transparent brown sticks. Structure 7OME depicts the azaCTZ-bound AncFT-L14 luciferase and 7QXQ the CEI-bound AncFT luciferase.

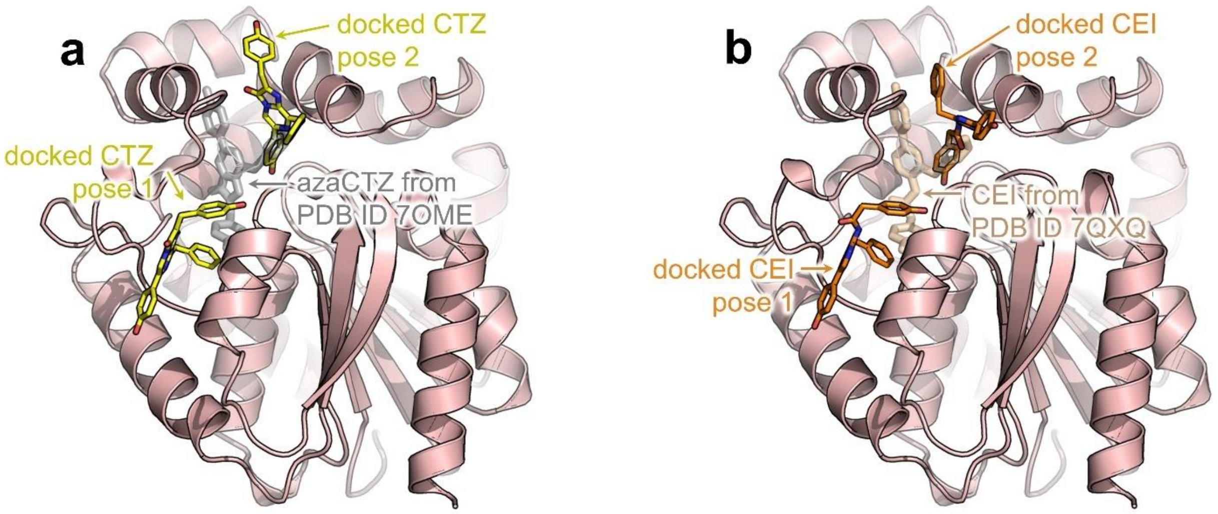

**Supplementary Fig. 11. Molecular docking of coelenterazine (CTZ) and coelenteramide (CEI) to the structure of Anc$^{HLD\text{-}RLuc}$ (PDB ID: 6G75) shown as a light red cartoon.** For both ligands, two energetically equivalent binding poses were predicted. (**a**) The predicted poses of CTZ shown as yellow sticks, and the crystallographic pose of azacoelenterazine (azaCTZ) as transparent grey sticks. (**b**) The predicted poses of CEI shown as orange sticks, and the crystallographic pose as transparent brown sticks. Structure 7OME depicts the azaCTZ-bound AncFT-L14 luciferase and 7QXQ the CEI-bound AncFT luciferase.

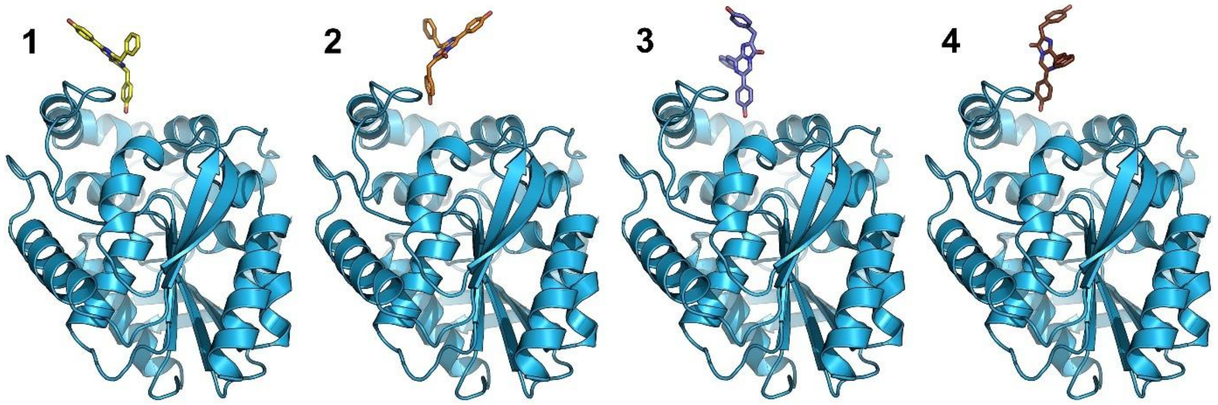

**Supplementary Fig. 12. Starting poses of coelenterazine (CTZ) for the simulation of substrate binding.** CTZ poses 1-4 are shown as yellow, orange, violet, and brown sticks, respectively. RLuc8 is shown as a blue cartoon.

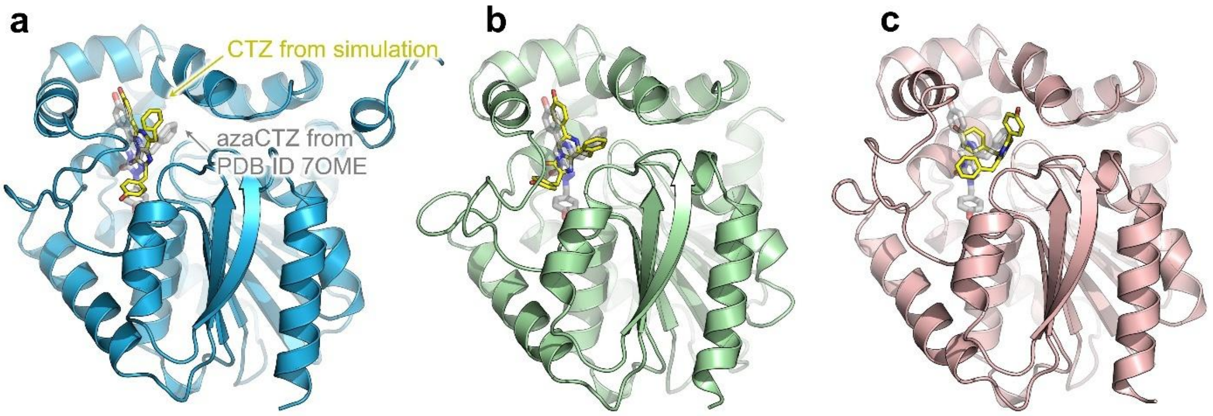

**Supplementary Fig. 13. Bound coelenterazine (CTZ, yellow sticks) from binding simulations started from pose 1 (Supplementary Fig. 12) using the ASMD method**. For reference, the crystallographic azacoelenterazine (azaCTZ) from the AncFT-L14 crystal structure (PDB ID: 7OME) is shown as transparent grey sticks. The binding was simulated in the RLuc8 structure (**a**; blue cartoon), AncFT-L14 (**b**; light green), and Anc$^{HLD\text{-}RLuc}$ (**c**; light red), PDB IDs: 2PSF, 7OME, and 6G75, respectively.

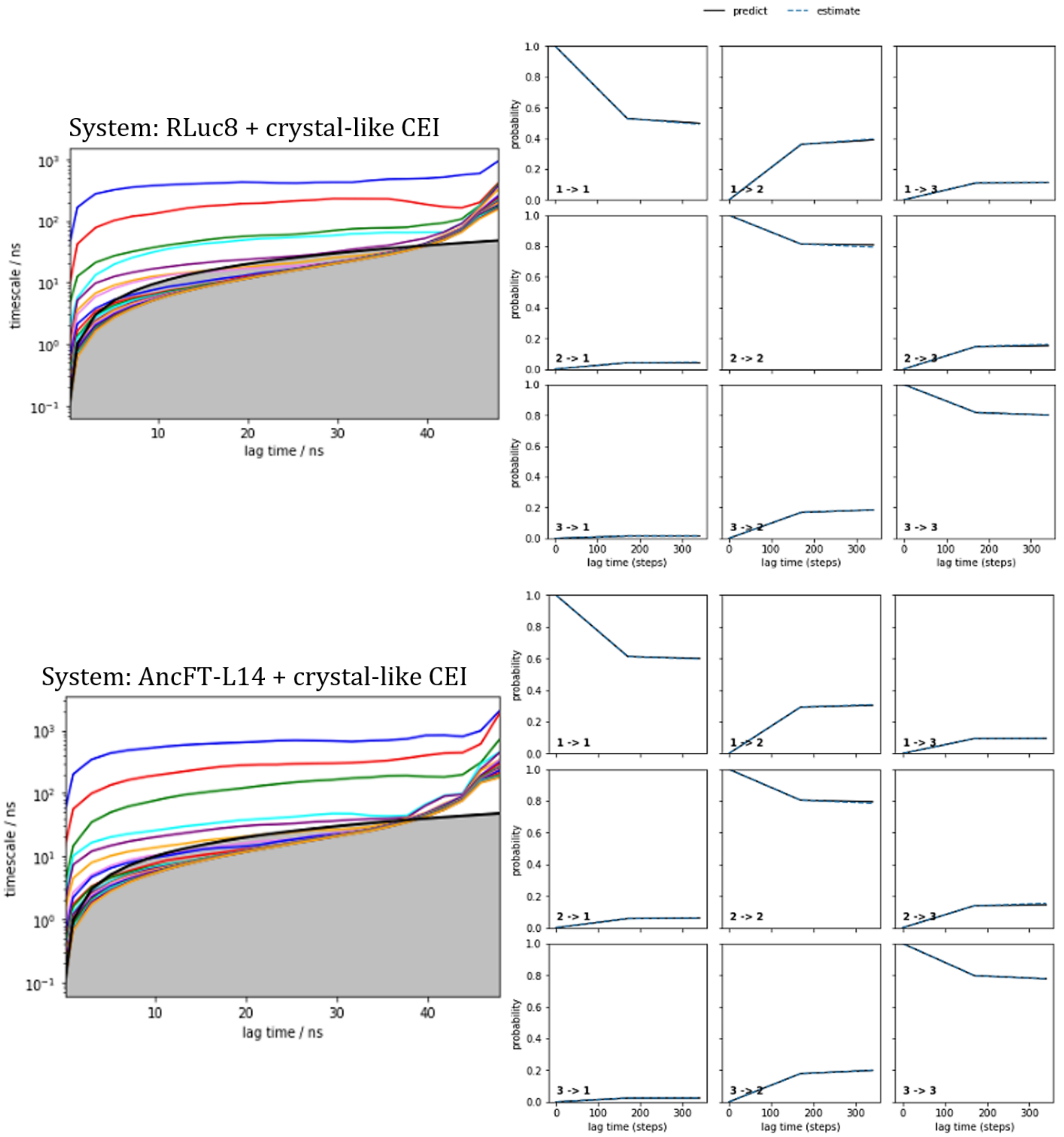

**Supplementary Fig. 14. Implied timescales and Chapman-Kolmogorov test of Markov state models (MSMs) of coelenteramide (CEI) release from RLuc8 and AncFT-L14.** The top row contains the results of CEI unbinding from RLuc8, while the bottom row from AncFT-L14. The implied timescales (left) show the transitions between the macrostates. The Chapman-Kolmogorov test (right) was done for an MSM of three macrostates constructed at a 17 ns lag time. The selected lag time is the lowest at which the “predict” and “estimate” lines agree.

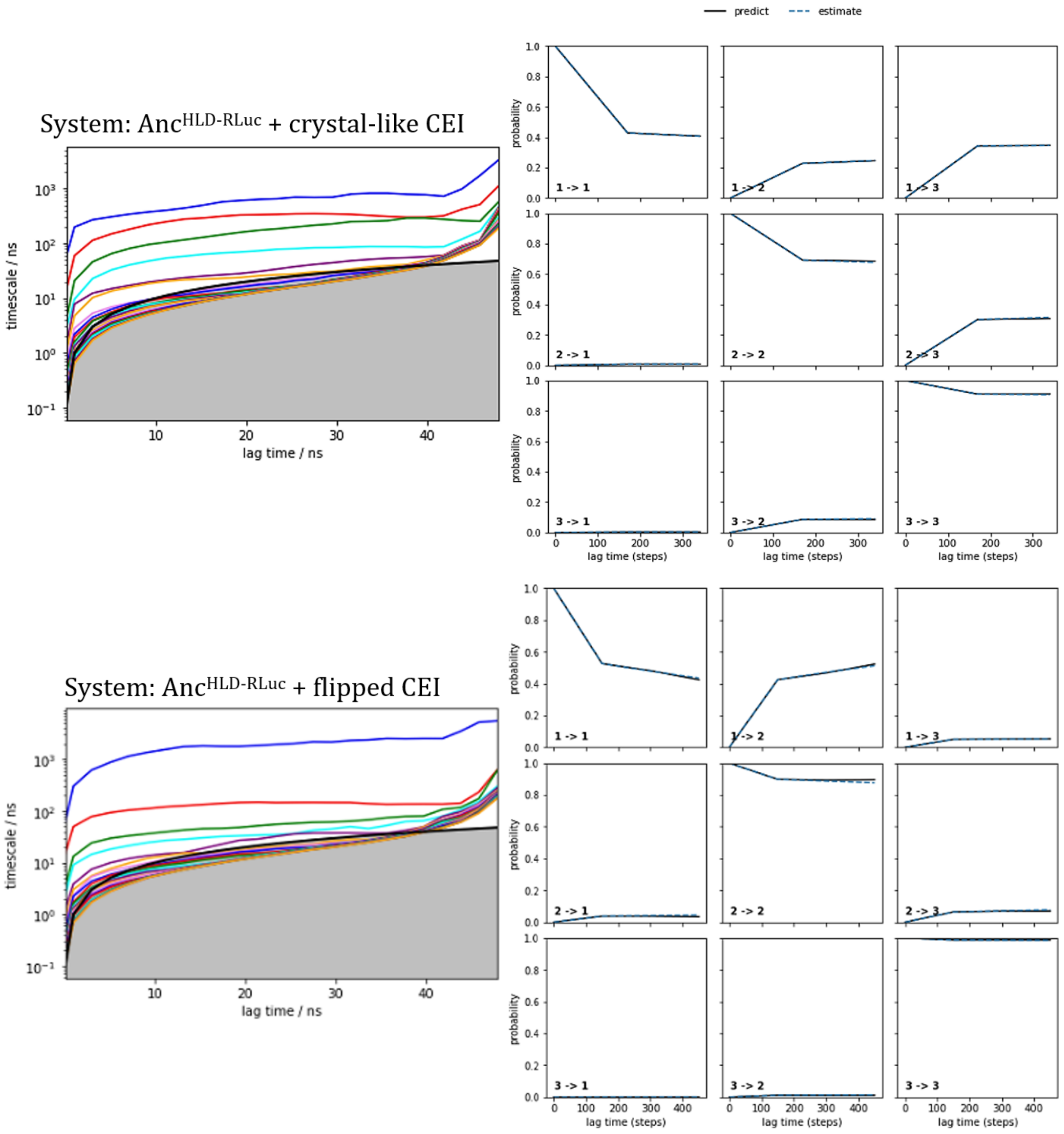

**Supplementary Fig. 15. Implied timescales and Chapman-Kolmogorov tests of Markov state models (MSMs) of coelenteramide (CEI) release from Anc$^{HLD\text{-}RLuc}$.** The top row contains the results of CEI unbinding from the crystal-like binding pose, while the bottom row corresponds to CEI release from the flipped pose. The implied timescales (left) show the transitions between the macrostates. The Chapman-Kolmogorov test (right) was done for an MSM of three macrostates for crystal-like CEI and four macrostates for flipped CEI. The MSM was constructed at a 17 ns lag time for crystal-like CEI and 15 ns for flipped CEI. The selected lag times are the lowest at which the “predict” and “estimate” lines agree.

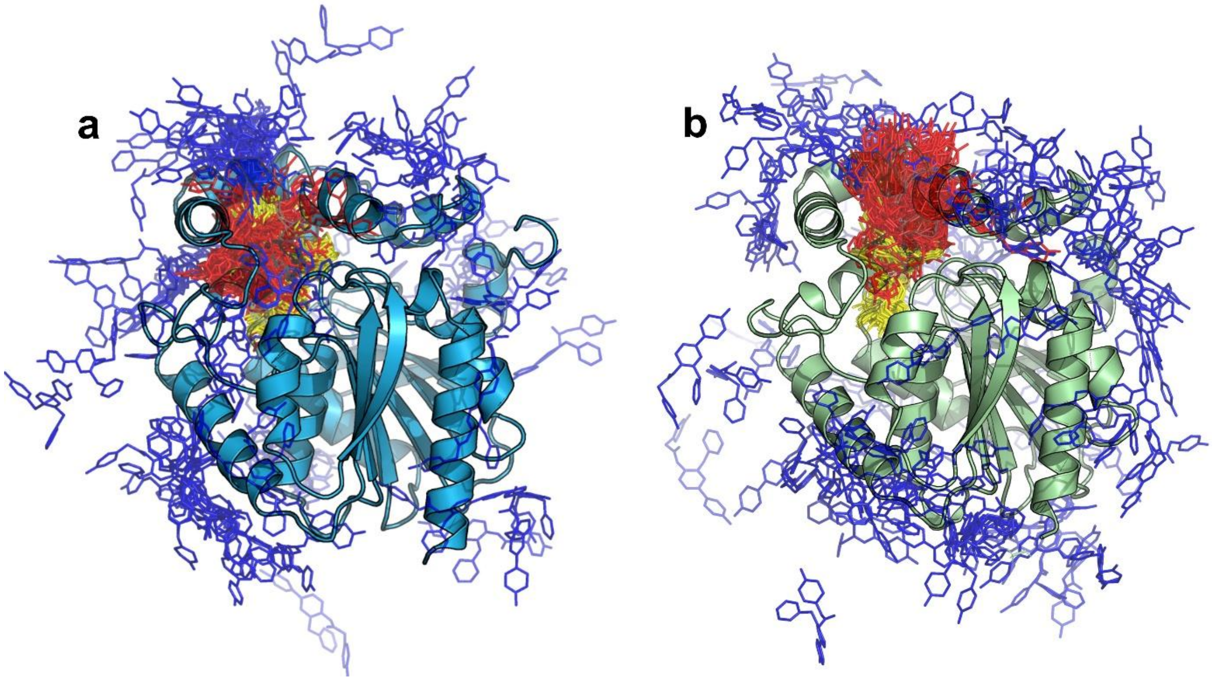

**Supplementary Fig. 16. Coelenteramide (CEI) macrostates from Markov state models (MSMs) from adaptive sampling of CEI unbinding from RLuc8 (a; blue cartoon) and AncFT-L14 (b; light green).** The macrostates are shown as thin sticks colored based on their distance to the active site (yellow, red, and blue in ascending order of distance).

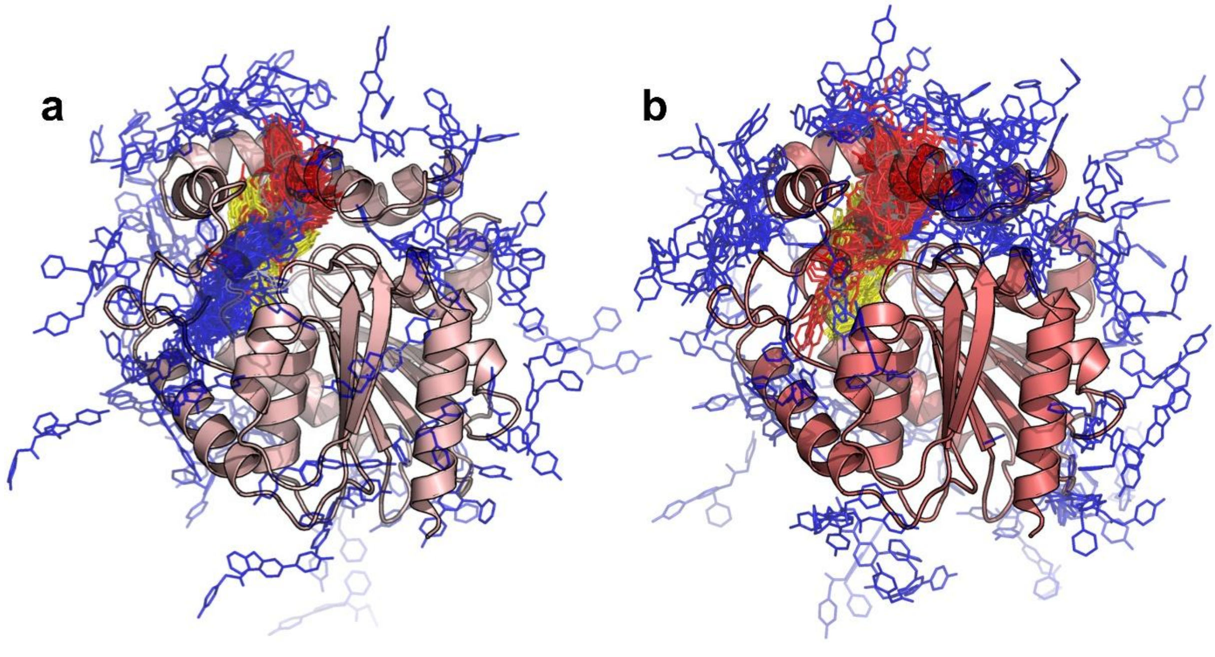

**Supplementary Fig. 17. Coelenteramide (CEI) macrostates from Markov state models (MSMs) from adaptive sampling of CEI unbinding from Anc$^{HLD-RLuc}$ (red cartoon).** Unbinding was simulated from the crystal-like CEI pose (**a**; light red protein structure) and the flipped CEI pose (**b**; dark red protein structure). The macrostates are shown as thin sticks colored based on their distance to the active site (yellow, red, and blue in ascending order of distance).

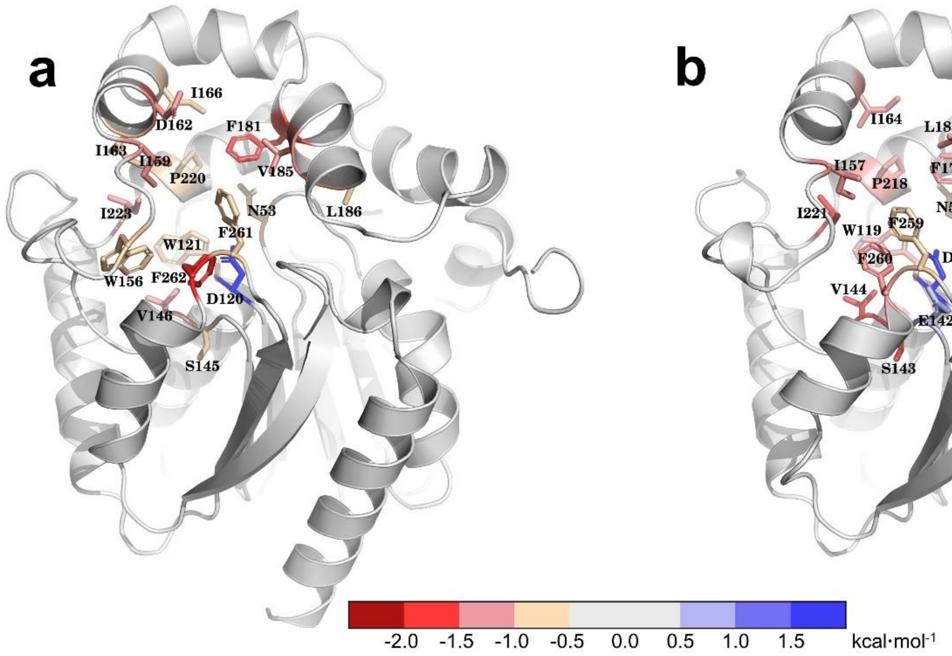

**Supplementary Fig. 18. Interactions of the 'bound' state of coelenteramide (CEI) in RLuc8 (a) and AncFT-L14 (b).** The interacting residues (<-0.5 and >0.5 $kcal.mol^{-1}$) are shown as sticks colored according to their interaction energy. All residue numbering corresponds to the numbering in RLuc8.

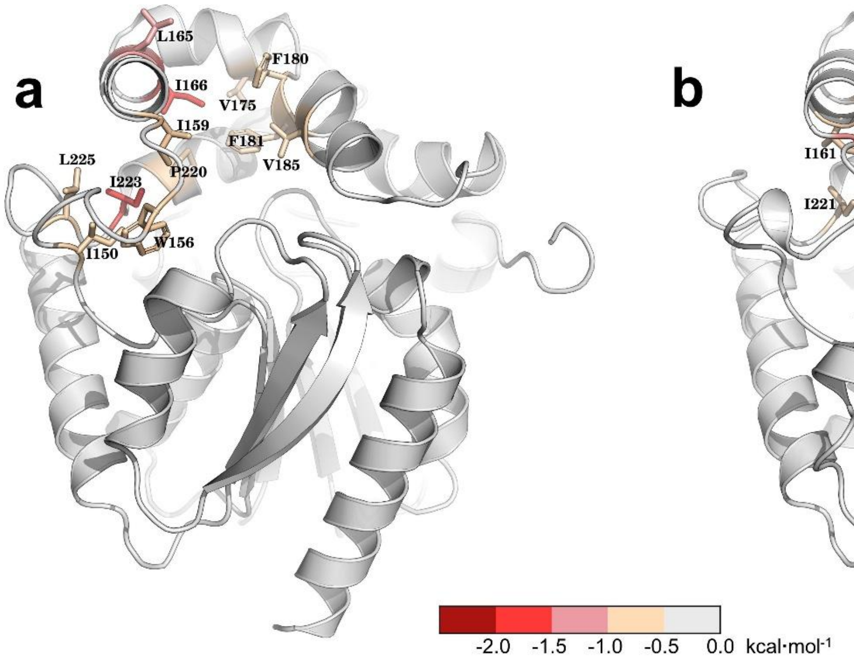

**Supplementary Fig. 19. Interactions of the 'tunnel' state of coelenteramide (CEI) in RLuc8 (a) and AncFT-L14 (b).** The interacting residues are shown as sticks colored according to their interaction energy. All residue numbering corresponds to the numbering in RLuc8.

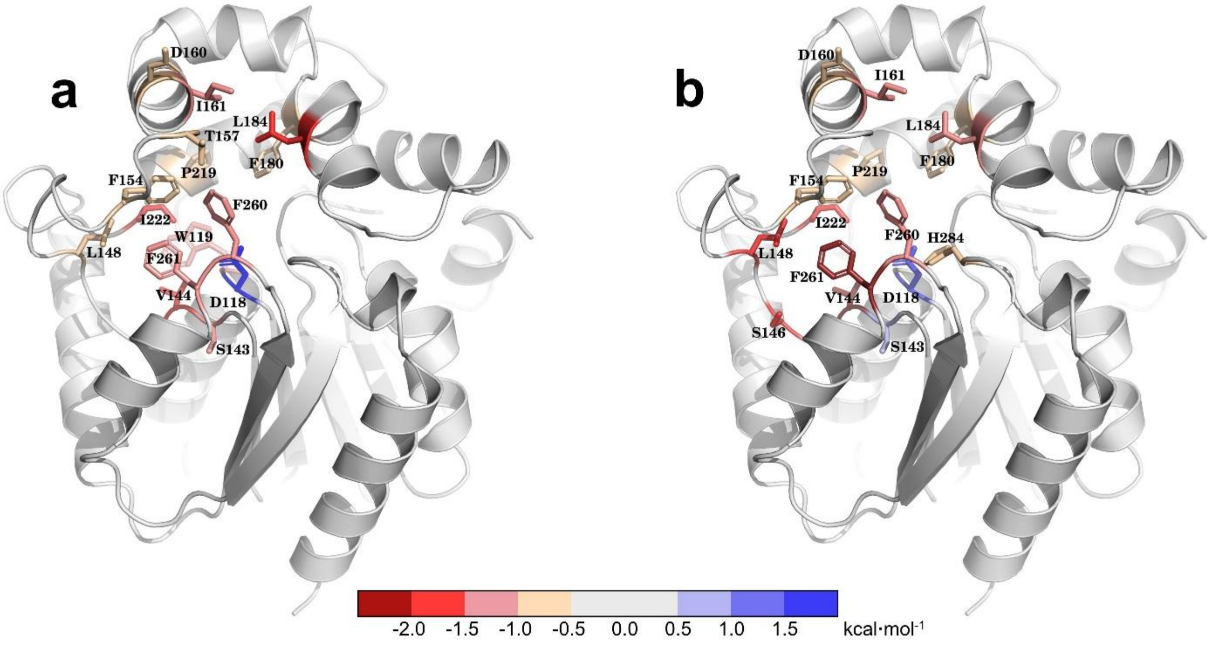

**Supplementary Fig. 20. Interactions of Anc$^{HLD-RLuc}$ with the 'bound' state of crystal-like coelenteramide (CEI) (a) and flipped CEI (b).** The interacting residues are shown as sticks colored according to their interaction energy. All residue numbering corresponds to the numbering in RLuc8.

### Supplementary Tables

**Supplementary Table I.** Values, standard errors, and confidence intervals of RLuc8 steady-state and inactivation kinetic parameters refined in this study and their comparison with values reported in the scientific literature. n.d. = not determined

| Parameter | Value | Standard error | Confidence intervals | Values in literature(*2-6*) |
|---|---|---|---|---|
| $K_{m,CTZ}$ [μM] | 2.16 | 0.02 | 2.02-2.28 | 1.3-2.9 |
| $K_{m,O2}$ [μM] | 719 | 6 | 669-780 | n.d. |
| $k_{cat}$ [$s^{-1}$] | 21.9 | 0.1 | 20.5-23.6 | n.d. |
| $k_{cat,app}$ [$s^{-1}$] | 5.65 | 0.03 | 5.29-6.09 | 4.5-5.1 |
| $K_p$ [μM] | 1.74 | 0.08 | 1.27-3.41 | 1.0-1.4 |
| $k_{inact}$ [$s^{-1}$] | 0.015 | 0.003 | 0.011-0.020 | n.d. |

**Supplementary Table II**. Values, standard errors, and confidence intervals of parameters derived from the pre-steady-state transient kinetic data of the *Renilla luciferase* RLuc8 catalysis. The parameters and their statistics were obtained by a global fitting of full-progress curves displayed in Figure 3 in the main text using numerical integration and by a confidence contour analysis with a $x^2$ threshold of 0.95. n.a. = not applicable

| Parameter | Value | Standard error | Lower bound | Upper bound |
|---|---|---|---|---|
| | | SUBSTRATES BINDING PROCESS | | |
| $k_{+1}$ [μM$^{-1}$.s$^{-1}$] | 1.76 | 0.05 | 1.55 | 2.16 |
| $E_{a,+1}$ [kcal.mol$^{-1}$] | 6.9 | 0.2 | 2.8 | 12.2 |
| $k_{-1}$ [s$^{-1}$] | 279 | 3 | 235 | 356 |
| $E_{a,-1}$ [kcal.mol$^{-1}$] | 6.7 | 0.2 | 1.1 | 12.1 |
| $k_{+2}$ [s$^{-1}$] | 34.9 | 1.4 | 25.1 | 46.6 |
| $E_{a,+2}$ [kcal.mol$^{-1}$] | 8.8 | 0.2 | 6.0 | 10.9 |
| $k_{-2}$ [s$^{-1}$] | 21.1 | 0.2 | 17.0 | 26.6 |
| $E_{a,-2}$ [kcal.mol$^{-1}$] | 9.2 | 0.1 | 7.9 | 12.8 |
| $K_{d,O_2}$ [μM] | 223 | 12 | 53 | 615 |
| $E_{a,O_2}$ [kcal.mol$^{-1}$] | 9.3 | 0.1 | 3.9 | 20.5 |
| | | CHEMICAL CONVERSION PROCESS | | |
| $k_{+3}$ [s$^{-1}$] | 0.244 | 0.006 | 0.037 | 0.512 |
| $E_{a,+3}$ [kcal.mol$^{-1}$] | 41 | 1 | 34 | 63 |
| | | PRODUCT RELEASE PROCESS | | |
| $k_{+4}$ [s$^{-1}$] | 0.16 | 0.03 | 0.08 | 0.48 |
| $E_{a,+4}$ [kcal.mol$^{-1}$] | > 19 | n.a. | 19 | n.a. |
| $k_{-4}$ [s$^{-1}$] | 1.2 | 0.2 | 0.6 | 1.6 |
| $E_{a,-4}$ [kcal.mol$^{-1}$] | < 22 | n.a. | 0 | 22 |
| $k_{+5}$ [s$^{-1}$] | 430 | 3 | 298 | 672 |
| $E_{a,+5}$ [kcal.mol$^{-1}$] | < 5.1 | n.a. | 0 | 5.1 |
| $k_{-5}$ [μM$^{-1}$.s$^{-1}$] | 1.77 | 0.01 | 1.41 | 2.22 |
| $E_{a,-5}$ [kcal.mol$^{-1}$] | 10 | 2 | 6 | 13 |

**Supplementary Table III**. Values, standard errors (S.E.), and confidence intervals of rate constants and activation energies obtained by global analysis of temperature-dependent transient kinetic data applying the induced-fit binding mechanism (AncFT, AncFT-L14) or simple one-step binding mechanism ($Anc^{HLD\text{-}RLuc}$). n.d. = not determined (not defined well by the data); n.a. = not applicable (no second step for the simple one-step binding mechanism)

| **Parameter** | **$Anc^{HLD\text{-}RLuc}$** | | **AncFT** | | **AncFT-L14** | |
|---|---|---|---|---|---|---|
| | Value ± S.E. | Confidence intervals | Value ± S.E. | Confidence intervals | Value ± S.E. | Confidence intervals |
| $k_{+1}$ [µM$^{-1}$.s$^{-1}$] | 0.033 ± 0.003 | 0.028-0.039 | 4.43 ± 0.06 | 4.21-4.71 | 4.52 ± 0.07 | 4.17-4.94 |
| $k_{-1}$ [s$^{-1}$] | 25 ± 4 | 19-35 | 258 ± 8 | 230-298 | 332 ± 10 | 283-394 |
| $k_{+2}$ [s$^{-1}$] | n.a. | n.a. | 19 ± 3 | 10-32 | 45 ± 3 | 32-66 |
| $k_{-2}$ [s$^{-1}$] | n.a. | n.a. | 25 ± 6 | 3-53 | 20 ± 3 | 10-34 |
| $E_{a,+1}$ [kJ.mol$^{-1}$] | 77 ± 4 | 71-84 | n.d. | n.d. | n.d. | n.d. |
| $E_{a,-1}$ [kJ.mol$^{-1}$] | 27 ± 7 | 17-36 | 51 ± 2 | 45-59 | 11 ± 2 | 3-22 |
| $E_{a,+2}$ [kJ.mol$^{-1}$] | n.a. | n.a. | < 59 | 0-59 | 94 ± 6 | 60-131 |
| $E_{a,-2}$ [kJ.mol$^{-1}$] | n.a. | n.a. | < 137 | 0-137 | 150 ± 12 | 120-187 |

**Supplementary Table IV.** Crystallographic data collection and refinement statistics for the determined co-crystal structure of AncFT-L14 complexed with azacoelenterazine (azaCTZ).

| **Data collection**[a] | **AncFT-L14/azaCTZ complex** |
|---|---|
| Wavelength [Å] | 0.999 |
| Resolution range [Å] | 39.44–1.501 (1.555–1.501) |
| Space group | $P2_12_12_1$ |
| Unit cell parameters a, b, c [Å] | 43.076, 64.487, 98.096 |
| Unit cell parameters α, β, γ [°] | 90, 90, 90 |
| Total reflections | 508,537 (26,381) |
| Unique reflections | 44,231 (4,130) |
| Multiplicity | 11.5 (6.4) |
| Completeness [%] | 99.34 (93.66) |
| Mean I/σ [I] | 15.33 (1.47) |
| Wilson B-factor [$Å^2$] | 12.16 |
| R-merge | 0.185 (1.228) |
| CC1/2 | 0.998 (0.619) |
| Reflections used in refinement | 44,231 (4,107) |
| Reflections used for R-free | 2,130 (191) |
| R-work [%] | 16.82 (25.9) |
| R-free [%] | 19.02 (27.33) |
| Number of non-hydrogen atoms | 2,665 |
| Macromolecules | 2,422 |
| Ligands | 32 |
| Solvent | 211 |
| Protein residues | 295 |
| RMS (bonds) [Å] | 0.006 |
| RMS (angles) [°] | 0.88 |
| Ramachandran favored [%] | 96.59 |
| Ramachandran allowed [%] | 3.41 |
| Ramachandran outliers [%] | 0.00 |
| Rotamer outliers [%] | 1.12 |
| Clash score | 2.91 |
| Average B-factor [$Å^2$] | 18.72 |
| Macromolecules | 17.84 |
| Ligands | 20.67 |
| Solvent | 28.53 |
| PDB ID | 7OME |

[a] Statistics for the highest-resolution shell are shown in parentheses.

**Supplementary Table V**. Predicted affinities of coelenterazine (CTZ) and coelenteramide (CEI) from molecular docking to RLuc8, AncFT-L14, and Anc$^{HLD\text{-}RLuc}$ (PDB IDs: 2PSF, 7OME, and 6G75, respectively).

| Protein | Ligand affinity (kcal.mol$^{-1}$) | |
|---|---|---|
| | CTZ | CEI |
| **RLuc8** | -12.1 | -11.3 |
| **AncFT-L14** | -13.2 | -12.2 |
| **Anc$^{HLD\text{-}RLuc}$** | -9.3 | -9.3 |

**Supplementary Table VI**. Summary of the mean coelenteramide (CEI)-active site distance and the equilibrium probabilities of macrostates from Markov state models from the adaptive sampling of CEI release from RLuc8, AncFT-L14, and Anc$^{HLD\text{-}RLuc}$.

| System | Macrostate | Mean distance (Å) | Equilibrium probability |
|---|---|---|---|
| **RLuc8 (crystal-like CEI)** | 'bound' | 5.17 ± 0.16$^{a}$ | 0.066 ± 0.028 |
| | 'tunnel' | 10.79 ± 0.34 | 0.506 ± 0.079 |
| | 'unbound' | 28.16 ± 0.31 | 0.428 ± 0.088 |
| **AncFT-L14 (crystal-like CEI)** | 'bound' | 4.59 ± 0.38 | 0.111 ± 0.052 |
| | 'tunnel' | 11.22 ± 0.87 | 0.485 ± 0.085 |
| | 'unbound' | 28.22 ± 0.50 | 0.404 ± 0.096 |
| **Anc$^{HLD\text{-}RLuc}$ (crystal-like CEI)** | 'bound' | 4.81 ± 0.07 | 0.009 ± 0.004 |
| | 'tunnel' | 9.99 ± 0.11 | 0.232 ± 0.066 |
| | 'unbound' | 25.00 ± 0.30 | 0.759 ± 0.067 |
| **Anc$^{HLD\text{-}RLuc}$ (flipped CEI)** | 'bound' | 6.11 ± 0.07 | 0.834 ± 0.063 |
| | 'tunnel' | 14.43 ± 0.23 | 0.153 ± 0.058 |
| | 'unbound' | 29.66 ± 0.46 | 0.014 ± 0.006 |

$^{a}$ The values were obtained by bootstrapping a random 80 % of the data 100 times.

**Supplementary Table VII**. Coelenteramide (CEI) unbinding kinetics from adaptive sampling simulations. The kinetic parameters were calculated from the Markov state models.

| System | RLuc8$^{e}$ (crystal-like CEI) | AncFT-L14 (crystal-like CEI) | Anc$^{HLD\text{-}RLuc}$ (crystal-like CEI) | Anc$^{HLD\text{-}RLuc}$ (flipped CEI) |
|---|---|---|---|---|
| **$\Delta G$ [kcal.mol$^{-1}$]$^{a}$** | 1.15 ± 0.31 | 0.81 ± 0.36 | 2.72 ± 0.34 | -2.33 ± 0.40 |
| **$k_{on}$ [M$^{-1}$.s$^{-1}$]$^{b}$** | (2.68 ± 0.80)×10$^{5}$ | (2.17 ± 0.73)×10$^{5}$ | (2.13 ± 0.84)×10$^{4}$ | (4.50 ± 0.91)×10$^{5}$ |
| **$k_{off}$ [s$^{-1}$]$^{c}$** | (1.14 ± 0.22)×10$^{6}$ | (7.09 ± 1.74)×10$^{5}$ | (1.26 ± 0.19)×10$^{6}$ | (3.86 ± 1.38)×10$^{4}$ |
| **$k_{off}/k_{on}$ [M]** | 5.1 ± 3.9 | 3.9 ± 2.1 | 80 ± 140 | 0.09 ± 0.05 |
| **$K_d$ [M]$^{d}$** | 8.2 ± 5.9 | 4.6 ± 2.6 | 140 ± 310 | 0.02 ± 0.01 |

$^{a}$ $\Delta G$ = the energy of the 'bound' state

$^{b}$ $k_{on}$ = rate constant of binding

$^{c}$ $k_{off}$ = rate constant of unbinding

$^{d}$ $K_d$ = dissociation constant of the 'bound' state

$^{e}$ The values were obtained from bootstrapping a random 80 % of the data 100 times.

### Supplementary Notes

**Supplementary Note 1. Global analysis of RLuc8 pre-steady-state transient kinetic data.** All the kinetic data (**Fig. 3** in the main text) were analyzed globally using the KinTek Explorer 10 software (KinTek, USA) which allows multiple data sets to be fit simultaneously to a single model(*7*). The fitting was based on the numerical integration of rate equations from the input model and experiment output observables, followed by applying the Bulirsch–Stoer algorithm with an adaptive step size to search a combination of parameters that produces the minimum $x^2$ value calculated from the nonlinear regression based on the Levenberg-Marquardt method. Residuals were normalized by sigma value for each data point. Performing the fit with the output observables corresponding to the experimental setup provided a very good fit of all the data, well-defining all the rate constants of the postulated mechanism. The luminescence data were first transformed to the cumulative luminescence data by integration and subsequently fit using **Equation 1** where S.O. is the experimental signal output, E.P and $E^*$.P is the enzyme-product complex in two different conformations, P is the reaction product, and $lum_T$ corresponds to the luminescence scaling factor defining the relationship between the luminescence signal output and product concentration at the temperature of the experiment (T).

$$\mathrm{S.O.} = lum_{\mathrm{T}} \cdot ([\mathrm{E.P}] + [\mathrm{E^*.P}] + [\mathrm{P}]) \tag{1}$$

The tryptophan fluorescence data were fit using **Equation 2** where S.O. is the experimental signal output, E is the enzyme, E.S and $E^*$.S is the enzyme-substrate (coelenterazine) complex in two different conformations, E.$O_2$ is the enzyme-oxygen complex, E.S.$O_2$ and E*.S.$O_2$ is the ternary enzyme-substrate-oxygen complex in two different conformations, E.P and $E^*$.P is the enzyme-product complex in two different conformations, $f_{E,T}$ is the fluorescence scaling factor defining the relationship between the fluorescence signal output and concentration of individual enzyme (E) species at the temperature of the experiment (T), and parameters $a_E$, $b_E$, $c_E$, and $d_E$ correspond to scaling factors defining ratios of fluorescence contributions of individual enzyme (E) species to the overall fluorescence signal.

$$\mathrm{S.O.} = f_{\mathrm{E,T}} \cdot \{([\mathrm{E}] + [\mathrm{E.O_2}]) + a_{\mathrm{E}} \cdot ([\mathrm{E.S}] + [\mathrm{E.S.O_2}]) + b_{\mathrm{E}} \cdot ([\mathrm{E^*.S}] + [\mathrm{E^*.S.O_2}]) + c_{\mathrm{E}} \cdot [\mathrm{E^*.P}] + d_{\mathrm{E}} \cdot [\mathrm{E.P}]\} \tag{2}$$

The coelenterazine fluorescence data were fit using **Equation 3** where S.O. is the experimental signal output, S is the coelenterazine substrate, E.S and $E^*$.S is the enzyme-substrate (coelenterazine) complex in two different conformations, E.S.$O_2$ and E*.S.$O_2$ is the ternary enzyme-substrate-oxygen complex in two different conformations, $f_{CTZ,T}$ is the fluorescence scaling factor defining the relationship between the fluorescence signal output and concentration of individual substrate (CTZ) species at the temperature of the experiment (T), and parameters $a_{CTZ}$ and $b_{CTZ}$ correspond to scaling factors defining ratios of fluorescence contributions of individual substrate (CTZ) species to the overall fluorescence signal.

$$\mathrm{S.O.} = f_{\mathrm{CTZ,T}} \cdot \{[\mathrm{S}] + a_{\mathrm{CTZ}} \cdot ([\mathrm{E.S}] + [\mathrm{E.S.O_2}]) + b_{\mathrm{CTZ}} \cdot ([\mathrm{E^*.S}] + [\mathrm{E^*.S.O_2}])\} \tag{3}$$

The coelenteramide fluorescence data were fit using **Equation 4** where S.O. is the experimental signal output, P is the coelenteramide product, E.P and $E^*$.P is the enzyme-product (coelenteramide) complex in two different conformations, $f_{CEI,T}$ is the fluorescence scaling factor defining the relationship between the fluorescence signal output and concentration of individual product (CEI) species at the temperature of experiment (T), and parameters $a_{CEI}$ and $b_{CEI}$ correspond to scaling factors defining ratios of fluorescence contributions of individual product (CEI) species to the overall fluorescence signal.

$$\mathrm{S.O.} = f_{\mathrm{CEI,T}} \cdot ([\mathrm{P}] + a_{\mathrm{CEI}} \cdot [\mathrm{E.P}] + b_{\mathrm{CEI}} \cdot [\mathrm{E^*.P}]) \tag{4}$$

The time-resolved absorbance spectra were first deconvoluted to individual components using singular value decomposition (SVD). The resulting time (amplitude) vectors were fit using **Equation 5** where $SVD_y$ is the reconstituted time dependence of the deconvoluted vector *y*, S is the coelenterazine substrate, P is the coelenteramide product, E is the enzyme, E.S and $E^*$.S is the enzyme-substrate (coelenterazine) complex in two different conformations, E.S.$O_2$ and E*.S.$O_2$ is the ternary enzyme-substrate-oxygen complex in two different conformations, E.P and $E^*$.P is the enzyme-product (coelenteramide) complex in two different conformations, and $svd_{xy}$ is the scaling factor defining the relationship between the SVD output and concentration of individual species. This notion assumed the presence of 5 species with different absorbance spectra in the measured data – (i) free substrate; enzyme-bound substrate; (iii) free product; (iv) enzyme-bound product; and (v) enzyme in any of the forms within the catalytic cycle. The change of the substrate and product absorbance spectra upon binding by enzyme was defined in accordance with previous experiments confirming spectral shifts upon binding(*3*).

$$\begin{aligned} \mathrm{SVD_y} &= svd_{\mathrm{xy}} \cdot [\mathrm{S}] + svd_{\mathrm{(x+1)y}} \cdot ([\mathrm{E.S}] + [\mathrm{E.S.O_2}] + [\mathrm{E^*.S}] + [\mathrm{E^*.S.O_2}]) + svd_{\mathrm{(x+2)y}} \cdot [\mathrm{P}] + svd_{\mathrm{(x+3)y}} \cdot ([\mathrm{E.P}] + [\mathrm{E^*.P}]) \\ &+ svd_{\mathrm{(x+4)y}} \cdot ([\mathrm{E}] + [\mathrm{E.O_2}] + [\mathrm{E.S}] + [\mathrm{E.S.O_2}] + [\mathrm{E^*.S}] + [\mathrm{E^*.S.O_2}] + [\mathrm{E.P}] + [\mathrm{E^*.P}]) \end{aligned} \quad (5)$$

The final global fit of the whole dataset combining all the aforementioned experimental data resulted in an unambiguous definition of all the rate constants with their unique constraints and no mutual correlations, as verified by the FitSpace Explorer(*8*) (**Supplementary Fig. *2***). In this analysis, each derived parameter was held fixed at various values while the other constants were allowed to float to achieve the minimal value of $x^2$. If $x^2$ increases as the fixed parameter diverges from the optimal value obtained from the fit, the constant is well-defined. By setting a threshold of acceptable $x^2$ values ($\min(x^2)/x^2$ = 0.95), the lower and upper limits of each parameter were determined (**Supplementary Table II**). By obtaining well-constrained and well-defined kinetic parameters for each step of the proposed kinetic pathway, the validity of the mechanism and reported parameters was justified.

**Supplementary Note 2. Kinetic analysis of the induced-fit binding process by the characterized variants Anc$^{\text{HLD-RLuc}}$, AncFT, and AncFT-L14.** Measurement of the coelenterazine substrate fluorescence upon mixing with an enzyme provided an increase of the optical signal upon binding the substrate inside the hydrophobic active site of the enzyme (**Supplementary Fig. 5**). Two kinetic phases, corresponding to two steps of the induced-fit binding mechanism, were observed for all the enzyme variants except for Anc$^{\text{HLD-RLuc}}$ which exhibited only one kinetic phase corresponding to a simple one-step binding mechanism. Such an outcome was in good agreement with the previously published data(*4*). By applying a temperature gradient using the rapid-mixing microfluidic chip, it was possible to uniquely determine rate constants as well as activation energies of all elementary binding steps for a thorough comparison of kinetic and thermodynamic parameters. In the case of AncFT and AncFT-L14, it was necessary to cool down the reaction to kinetically capture the fast initial kinetic phase which was hidden in the dead-time when elevated temperatures were applied. In contrast, the very slow kinetics of Anc$^{\text{HLD-RLuc}}$ required heating to speed up the binding process and reliably monitor the kinetics of binding. Thanks to the very high melting temperature of Anc$^{\text{HLD-RLuc}}$, it was possible to heat up the microfluidic chip up to 49 °C without discriminating the enzyme activity. Furthermore, the simple binding mechanism of Anc$^{\text{HLD-RLuc}}$, exhibiting only one kinetic phase, made it possible to collect data at fewer temperature points while still reliably determining thermodynamic parameters, as validated by the confidence contour analysis (**Supplementary Fig. 6**). On the other hand, some of the steps of the induced-fit binding, detected for AncFT and AncFT-L14, showed insignificant changes with decreasing temperature, preventing constrained determination of all activation energy values. Specifically, the activation energy of the first forward step ($E_{a,+1}$) was not defined by the data at all for both AncFT and AncFT-L14. Additionally, only the upper limits of activation energies of the second step ($E_{a,+2}$, $E_{a,-2}$) could be reliably derived from the data in the case of AncFT (**Supplementary Fig. 6**). All these findings were assumed and reported accordingly to prevent overinterpretation of the collected data and their global analysis (**Supplementary Table III**).

**Supplementary Note 3. Crystallographic analysis of the azacoelenterazine (azaCTZ)-bound co-crystal structure of AncFT-L14.** AzaCTZ-bound AncFT-L14 structure shows a canonical αβα-sandwich architecture, similar to the previously reported AncFT structures(*3*), with root-mean-square deviation (RMSD) value on Cα atoms ranging from 0.4 Å (**Supplementary Fig. 7**). Inspection of the electron density map unambiguously revealed azaCTZ molecule bound in the active site of the enzyme, adopting a Y-shaped conformation that is characteristic for *Renilla*-type catalysis (**Fig. 7b** in the main text). We observe that the overall binding mode and molecular contacts of azaCTZ in AncFT-L14 are very similar, if not identical, to the previously characterized AncFT luciferase(*3*). Additionally, there are unambiguously resolved two water molecules in the azaCTZ-bound AncFT-L14 structure, positioned in the putative dioxygen-binding site (**Fig. 7b-c** in the main text). These two waters are attracted to each other through a hydrogen bond (2.7 Å), while a former water is further hydrogen bonded with a carboxylate of D118 (2.8 Å) and a side chain carbonyl of N51 (3.1 Å). The latter water then forms a hydrogen bond with a main chain amide nitrogen of N51 (3.0 Å). Due to this configuration, the former water molecule is positioned over the triazolopyrazine core of azaCTZ, in close proximity (3.5 Å) to its N1-nitrogen. In the native CTZ, the C2-carbon is present in this position, which is the site where the dioxygen is incorporated into the luciferin. The presence and localization of water molecules at the bottom of the active site pocket thus mimic, to a certain extent, the binding of dioxygen in a Michaelis enzyme-substrate complex. Our structural observations highlight that the side chain carboxylate of D118 can be involved, together with N51 and W119, in the positioning of a co-substrate molecule (dioxygen) such that it can be directly attacked by the C2 carbon of activated CTZ.

**Supplementary Note 4. Molecular docking and adaptive steered molecular dynamics (ASMD) analysis of ligand binding to active sites of Anc$^{HLD\text{-}RLuc}$, AncFT-L14, and RLuc8.** Coelenterazine (CTZ) substrate and coelenteramide (CEI) product were docked to RLuc8, AncFT-L14, and Anc$^{HLD\text{-}RLuc}$. The predicted poses resemble the crystallographic luciferins in structures PDB ID 7OME (AncFT-L14 + azaCTZ), 7QXR (AncFT + azaCTZ), and 7QXQ (AncFT + CEI), except for Anc$^{HLD\text{-}RLuc}$ (**Supplementary Figs. 9-11**), which also showed worse binding energies than the other enzymes (**Supplementary Table V**). This finding served as a basis for the follow-up substrate binding simulation by pulling CTZ into the active site using ASMD. We used four different orientations of CTZ at the mouth of the main access tunnel (**Supplementary Fig. 12**) as the starting points to obtain different binding poses and to evaluate the potential reactivity of the most probable pose. For RLuc8 and AncFT-L14, the binding was easiest from pose 1, which resembles the crystal-like orientation of luciferin in PDB ID 7OME. In contrast, Anc$^{HLD\text{-}RLuc}$ preferred starting pose 3, which is flipped compared to pose 1, the pose favored by RLuc8 and AncFT-L14. Consequently, the simulation resulted in a flipped binding pose (**Fig. 8b** in the main text), a conformation geometrically not well-prepared for the catalysis. Similarly, CTZ binding to Anc$^{HLD\text{-}RLuc}$ from pose 1 resulted in a poorly oriented binding mode with even worse energy than pose 3 (**Supplementary Fig. 13**).

**Supplementary Note 5. Product release analysis by adaptive sampling molecular dynamics and definition of macrostate clustering.** The release of the coelenteramide (CEI) product was simulated with adaptive sampling molecular dynamics starting from the crystal-like pose in each enzyme and additionally, from the flipped pose in Anc$^{HLD\text{-}RLuc}$. Note that the crystal-like pose of CEI in Anc$^{HLD\text{-}RLuc}$ was artificially prepared by transplanting the ligand from crystal structure PDB ID 7QXQ since this pose was not observed experimentally nor predicted by docking. The simulations were clustered to separate the 'bound' CEI state (CEI in the active site), the 'tunnel' state (CEI in the access tunnel), and the 'unbound' CEI state (**Supplementary Figs. 14-17**), to calculate their equilibrium probabilities (**Supplementary Table VI**), and to estimate kinetics (**Supplementary Table VII**). RLuc8 and AncFT-L14 simulations had comparable results, indicating that the product release is energetically favorable in both enzymes (positive $\Delta G$ of the 'bound' state, $k_{off}$ faster than $k_{on}$). In the 'bound' macrostates, CEI interacted mainly with the same residues, except for D162, which had a significant affinity with CEI only in RLuc8 (**Supplementary Fig. 18**). However, the 'tunnel' macrostates look significantly different (**Supplementary Fig. 16**). During the product release, the conformation of RLuc8 was changing. CEI was interacting with the dynamical L9/α4 fragment (**Supplementary Fig. 19**), which was moving to the side, making the access tunnel wider (**Fig. 8d** in the main text). In contrast, AncFT-L14 was much more rigid in this area, so CEI left the active site straight through the tunnel. Therefore, the per-residue interactions differ noticeably (**Supplementary Fig. 19**). In RLuc8, the interaction energy is considerably higher for W156, L165, I166, V175, and I223 (most in α4 and L9), while in AncFT-L14, I159, V185, and F261 are preferred. The simulations with Anc$^{HLD\text{-}RLuc}$ had very different results depending on the starting orientation of CEI. The release of CEI from Anc$^{HLD\text{-}RLuc}$ from the transplanted crystal-like pose followed the same trend as RLuc8 and AncFT-L14 but with a more pronounced difference in the probabilities of macrostates (**Supplementary Table VI**). With the ‘bound’ state 90 times less probable than the ‘unbound’, this system would likely dissociate rapidly. Moreover, CEI was not released through the main access tunnel but through a new opening between L9 and α9 elements, which probably formed as the artificial binding pose disrupted the enzyme. In contrast, the release of the flipped CEI (which corresponds to the preferred binding orientation of CTZ, as described above) followed the opposite trend, with the ‘bound’ state significantly more probable than the ‘unbound’ state (**Supplementary Table VI**). High stability of the enzyme-product complex could cause product inhibition, confirmed by previous kinetic analyses of $Anc^{HLD-RLuc}$(*4*). Indeed, CEI interacted with different residues in the ‘bound’ state depending on its orientation (**Supplementary Fig. 20**). For example, flipped CEI had a stronger affinity towards V146, S148, and L150 than the crystal-like pose. Regarding flexibility, $Anc^{HLD-RLuc}$ is the most rigid of the three enzymes (**Fig. 8g** in the main text). Moreover, it has the narrowest access tunnel (**Fig. 8d** in the main text), which explains the difficulty of ligand binding and unbinding (**Supplementary Tables V-VII**).

### Materials and Methods

*Luciferin and aza-luciferin molecules*

The coelenterazine (CTZ) luciferin was purchased from P-LAB (cat. no. R4094.3). Substrate analogue azacoelenterazine (azaCTZ) was synthesized as described previously(*3*).

*Loop transplantation to create AncFT-L14*

The mutant construct was generated using standard PCR-based nested protocols and inserted into the pET21b expression vector between NdeI and BamHI sites. Briefly, the L14 loop exchange mutant was designed based on sequence and structural comparison. The correlated motions of loop L9 and loop L14 were identified using the in-house web-based tool LoopGrafter(*9, 10*). The RLuc L14 loop sequence $L_{235}$VKGGKP was introduced instead of the corresponding element encompassing $I_{223}$KGDGPE sequence in AncFT(*4*) to create the AncFT-L14 mutant. Loop transplantation was carried out in two-step PCR using Phusion polymerase (NEB, UK) according to the manufacturer's protocol. The list of the used primers is available in **Table SVIII**. After the mutagenesis reaction, the original template was removed by DpnI (NEB, USA) treatment (2 h at 37°C), followed by the DpnI inactivation (20 min at 80°C). The resulting plasmids were transformed into chemocompetent *E. coli* DH5α cells, plated on LB-agar (tryptone 10 $g.L^{-1}$, yeast extract 5 $g.L^{-1}$, NaCl 10 $g.L^{-1}$, agar 15 $g.L^{-1}$) containing ampicillin (100 $μg.mL^{-1}$), and incubated at 37 °C overnight. Plasmids were isolated from three randomly selected colonies, and error-free sequences were verified by DNA sequencing (Eurofins Genomics, Germany).

**Supplementary Table VIII: Nucleotide primers used in PCR mutagenesis to generate AncFT-L14.**

| Primer name | Length (nt) | Nucleotide sequence (5' → 3') |
|---|---|---|
| FWD1 | 20 | TAATACGACTCACTATAGGG |
| RVS1 | 22 | CGGAATTTCACGAGGCCAGGTC |
| FWD2 | 67 | GACCTGGCCTCGTGAAATTCCGCTGGTTAAAGGTGGTAAACCGGATGTGATTGAAATTGTGAAAAGC |
| RVS2 | 19 | GCTAGTTATTGCTCAGCGG |

*Overproduction and purification of $Anc^{HLD-RLuc}$, AncFT, AncFT-L14, and RLuc8*

Overexpression of proteins from pET21b plasmid ($amp^R$) was carried out in *E. coli* BL21 cells (NEB, USA) cultivated in LB medium supplemented with ampicillin (100 $μg.mL^{-1}$). Once the culture $OD_{600}$ reached ~0.6, protein production was induced at 20 °C by adding IPTG to a final concentration of 0.5 mM. The cells were then harvested after 16 h incubation by centrifugation (4,000 rpm, 20 min, 4 °C), and the cell pellets were resuspended in 20 mM potassium phosphate buffer pH 7.5, containing 500 mM NaCl and 10 mM imidazole and stored at -80°C. Prior to purification, the thawed cell mass was disrupted by sonication using Sonic Dismembrator Model 705 (Fisher Scientific, USA). Lysates were clarified by centrifugation (14,000 rpm, 1 h, 4 °C) using a Sigma 6-16K centrifuge (SciQuip, UK) equipped with a 12166 rotor. Supernatants containing the recombinant His-tagged proteins at their C-terminal ends were metal-affinity purified using Ni-NTA Superflow Cartridge 5mL (Qiagen, Germany) installed on FPLC system (Bio-Rad Laboratories, USA) and equilibrated with the 20 mM potassium phosphate buffer pH 7.5, containing 500 mM NaCl and 10 mM imidazole. Proteins were eluted with an imidazole gradient and monomers were separated using gel permeation chromatography on Äkta FPLC (GE Healthcare, Sweden) equipped with HiLoad™ 16/600 Superdex™ 200 pg column (GE Healthcare, Sweden) and equilibrated with 10 mM Tris-HCl buffer pH 7.5 containing 50 mM NaCl. Purified proteins were concentrated to final concentrations using Centrifugal Filter Units Amicon® Ultra-15 Ultracel®-10K (Merck Millipore Ltd., Ireland). The purity of proteins was verified on SDS-PAGE. The concentration of protein samples was measured using DeNovix® DS-11 Spectrophotometer (DeNovix Inc., USA).

*Circular dichroism spectroscopy*

Circular dichroism spectra were recorded at 20 °C using the Chirascan spectropolarimeter (Applied Photophysics, UK). The signal was measured from 190 to 260 nm at a rate of 100 $nm.min^{-1}$, with 1.0 s integration time, and 1 nm bandwidth in a 0.1 cm quartz cuvette. The spectrum was measured in five consecutive technical replicates and then averaged and corrected for the ellipticity of the buffer. The circular dichroism signal was expressed as mean residue ellipticity $\Theta_{MRE}$, calculated using **Equation 6** where $\Theta_{obs}$ is the measured ellipticity in degrees, $M_W$ is the protein molecular weight in kDa, $n$ is the number of protein residues, $l$ is the cell path length in cm, and $c$ is the protein concentration in $mg.mL^{-1}$.

$$\Theta_{\mathrm{MRE}} = \frac{\Theta_{\mathrm{obs}} \cdot M_{\mathrm{W}} \cdot 100}{n \cdot c \cdot l} \tag{6}$$

*Thermal stability measurements*

Thermal unfolding was studied using NanoDSF Prometheus (NanoTemper, Germany) by monitoring tryptophan fluorescence over the temperature range of 20 to 90 °C, at a heating rate of 1 °C/min. The onset and melting temperatures ($T_{onset}$ and $T_m$, respectively) were evaluated directly by the manufacturer-provided software ThermControl v2.0.2.

*Bioluminescence emission spectral analysis*

Bioluminescence emission spectra were measured at ambient temperature using the spectrofluorometer FluoroMax-4 (HORIBA, Japan). 1 mL of an enzyme solution in 100 mM potassium phosphate buffer pH 7.5 was added into a quartz cuvette and the enzymatic reaction was initiated by manual injection of 50 µL of ethanolic solution of coelenterazine followed by a quick mixing. The acquisition of luminescence emission spectra started 10 seconds after the addition of the coelenterazine substrate. The resulting concentration of an enzyme in the reaction mixture ranged from 0.7 to 1.0 µM while the concentration of coelenterazine was approximately 30 µM. The data were collected for wavelengths from 350 to 700 nm with a step of 2 nm and each spectrum was measured in triplicates. The obtained spectrum was normalized relative to the value of the highest peak for each of the enzymes individually and the shape and positions of the luminescence emission peaks were compared.

*Bioluminescence emission decay and stability*

NIH/3T3 mouse fibroblast cells (ATCC® CRL-1658™) were transfected according to the manufacturer protocol using Lipofectamine 2000 (Thermo Fisher, USA) with pcDNA3.1(+) plasmids containing the genes of luciferases codon-optimized for expression in mammalian cells (Gene Art, Thermo Fisher, USA). Cells were lysed 24 hours after the transfection, and the bioluminescence signal in the lysate was measured with a microplate reader FLUOstar Omega (BMG Labtech, Germany) after the addition of coelenterazine at the final concentration of 4.5 µM. Cells transfected with an empty pcDNA3.1(+) plasmid were used as a negative control. The measurements were done in three independent replicates.

*Conventional steady-state kinetic experiments (physiological oxygen atmosphere)*

Solid coelenterazine was dissolved in ice-cold ethanol and stored under a nitrogen atmosphere in dark glass vials at –20 °C. Before each measurement, the concentration and maintained quality of the ethanol stock solutions were verified spectrophotometrically. A buffer solution of coelenterazine was prepared by mixing an appropriate volume of the ethanol stock solution with 100 mM phosphate buffer pH 7.5 immediately before the measurement. Tested luciferase enzyme samples were diluted in the same 100 mM phosphate buffer pH 7.5. The enzymatic reaction was initiated inside the microplate reader FLUOstar OPTIMA (BMG Labtech, Germany) by the automatic addition of 225 µL buffer solution of coelenterazine into a microplate well containing 25 µL of enzyme solution. The resulting bioluminescence activity traces were collected at 37 °C until the luminescence intensity decreased to less than 0.5 % of its maximal measured value to ensure the full conversion of coelenterazine. Each luminescence trace was measured in 3 repetitions. The concentration of coelenterazine in the final reaction mixture ranged from 0.2 to 7.9 μM while the concentration of luciferase enzymes ranged from 17 to 640 nM.

*Oxygen-dependent kinetic experiments*

100 mM potassium phosphate buffer pH 7.5 and water solutions were incubated in the anaerobic glovebox Belle MR2 (Belle Technology, UK) overnight while stirring to remove dissolved oxygen. The glovebox was filled with ultra-high purity (UHP) nitrogen atmosphere containing a residual oxygen concentration of 2.4 ppm, as determined by the oxygen meter O2M-3 (Belle Technology, UK). A bottle with lyophilized evacuated (oxygen-free) RLuc8 protein was put into the anaerobic glovebox as well, it was dissolved with anaerobic water and incubated for at least one hour to remove possible traces of dissolved oxygen. Finally, a nitrogen-flushed aliquot of CTZ dissolved in pure ethanol was put into the glovebox, incubated for approximately 20 minutes to remove dissolved oxygen, and then closed to prevent further evaporation of the sample. Aerobic buffer solutions of controlled oxygen concentrations were prepared by filling 100 mM potassium phosphate buffer pH 7.5 into glass tonometers and flushing/bubbling the solution with a gas mixture containing different ratios of oxygen and nitrogen for at least one hour. Afterward, the tonometers were hermetically closed, put into the anaerobic glovebox, and kept closed before loading into the Stopped-Flow instrument to prevent leakage of oxygen into the glovebox. Oxygen concentration in the tonometer solutions was calculated based on Henry's Law using known experimental temperature and pressure.

CTZ stock solution was diluted with 100 mM potassium phosphate buffer pH 7.5 and then manually premixed anaerobically with the RLuc8 enzyme to reach the desired concentration of both components. The premixed RLuc8.CTZ complex was loaded into the first port of the Stopped-Flow instrument KinetAsyst SF-61DX2 (TgK Scientific, UK) while the second one was filled with the aerobic buffer from the prepared tonometer. The two solutions were rapidly mixed in a 1:1 ratio and the enzymatic reaction was followed by an increase of the luminescence signal over 60 seconds at 37 °C. The port filling was repeated for 10 different CTZ concentrations and 5 different oxygen concentrations in a fully combinatorial manner (50 different conditions measured in total). The resulting concentrations during the experiments were 8 nM RLuc8, 0.2–15 μM CTZ, and 61–829 μM oxygen. By applying logarithmic distribution of data points collection (high frequency at the beginning and lower frequency at the end of the collection), both pre-steady-state and steady-state phases of the reaction could be accurately monitored.

*Steady-state kinetic data analysis*

The obtained kinetic curves were integrated in order to obtain changes of cumulative luminescence over time that are proportional to the product formation increase with time. The resulting curves were analyzed by applying an updated protocol employing global numerical fitting of raw kinetic data using KinTek Explorer 10 (KinTek Corporation, USA). The software allows for the input of a given kinetic model via a simple text description, and the program then derives the differential equations needed for numerical integration automatically. Numerical integration of rate equations searching a set of kinetic parameters that produce a minimum $x^2$ value was performed using the Bulirsch-Stoer algorithm with adaptive step size, and nonlinear regression to fit data was based on the Levenberg-Marquardt method. Residuals were normalized by sigma value for each data point. The standard error (S.E.) was calculated from the covariance matrix during nonlinear regression. In addition to S.E. values, a more rigorous analysis of the variation of the kinetic parameters was accomplished by confidence contour analysis using FitSpace Explorer (KinTek Corporation, USA). In these analyses, the lower and upper limits for each parameter were derived from the confidence contour obtained from setting $x^2$ threshold at 0.95. In the case of oxygen-dependent data, a general random order binding two-substrate steady-state model with irreversible enzyme inactivation (**Figure S21**) was used to obtain realistic values of turnover number $k_{cat}$, Michaelis constants $K_{m,CTZ}$ and $K_{m,O_2}$, product inhibition constant $K_p$, and inactivation rate constant $k_{inact}$ while in the case of 'conventional' (physiological oxygen atmosphere) experiments, a simpler minimal steady-state model (**Figure S22**) was used to derive $k_{cat}$, $K_{m,CTZ}$, and $K_p$. A conservative estimate for diffusion-limited substrates and product binding $k_{+1}$, $k_{+2}$, and $k_{-4}$ (100 μM$^{-1}$.s$^{-1}$) was used as a fixed value to mimic the rapid equilibrium assumption.

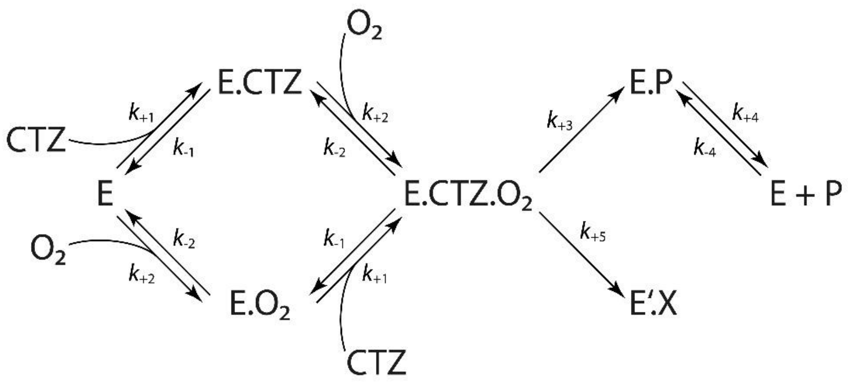

$$K_{m,CTZ} = k_{-1}/k_{+1} = k_{-1}/100 \qquad K_{m,O_2} = k_{-2}/k_{+2} = k_{-2}/100$$

$$k_{cat} = k_{+3} \qquad k_{inact} = k_{+5} \qquad K_p = k_{+4}/k_{-4} = k_{+4}/100$$

**Supplementary Fig. 21: Kinetic scheme of the random order two-substrate binding steady-state model.** Except for the turnover number $k_{cat}$ and Michaelis constant $K_m$ for both substrates, the model also includes an irreversible enzyme inhibition and product inhibition, described by $k_{inact}$ and $K_p$, respectively. Calculation of the steady-state kinetic parameters from the respective elementary rate constants is provided below the kinetic scheme.

$$E + CTZ \underset{k_{-1}}{\overset{k_{+1}}{\rightleftharpoons}} E.CTZ \xrightarrow{k_{+3}} E.P \underset{k_{-4}}{\overset{k_{+4}}{\rightleftharpoons}} E + P$$

$$K_{m,CTZ} = k_{-1}/k_{+1} = k_{-1}/100 \qquad k_{cat} = k_{+3} \qquad K_p = k_{+4}/k_{-4} = k_{+4}/100$$

**Supplementary Fig. 22: Kinetic scheme of the simple minimal steady-state model.** The model is capable of accurately determining values of the turnover number $k_{cat}$, Michaelis constant for the coelenterazine (CTZ) substrate $K_{m,CTZ}$, and product inhibition constant $K_p$. Calculation of the steady-state kinetic parameters from the respective elementary rate constants is provided below the kinetic scheme.

*Tryptophan fluorescence and luminescence stopped-flow kinetic experiments*

Transient kinetic traces upon the coelenterazine (CTZ) substrate conversion by the RLuc8 enzyme were monitored after rapidly mixing the components using the Stopped-Flow SFM 3000 mixing system equipped with a xenon arc lamp light source and the MOS-500 spectrometer (BioLogic, France). In each case, the reaction was initiated by mixing 75 µL of the enzyme solution with 75 µL of the CTZ solution with a total flow rate of 13 $mL.min^{-1}$ and then monitored in two modes using two different sources of the optical signal: (i) native tryptophan fluorescence and (ii) bioluminescence. Quenching of the tryptophan fluorescence, caused by the CTZ substrate binding in the active site, was monitored using the 340 ± 13 nm band-pass emission filter after the excitation at 295 nm. Bioluminescence signal changes, direct output of the luciferase reaction upon the substrate conversion, were monitored without applying any emission filter and no external excitation light source. The experiment was performed in 100 mM potassium phosphate buffer pH 7.5 at either 15 °C or 37 °C in three different experimental setups: (i) multiple turnover (excess of the substrate) varying the CTZ concentration; (ii) multiple turnover varying the RLuc8 concentration; and (iii) single turnover experiment (excess of the enzyme). Each kinetic trace was collected in 6 consecutive technical replicates and then averaged. The resulting concentrations after mixing were 2 µM RLuc8 and 1.3–64 µM CTZ in the CTZ-varying multiple turnover experiment, 15 µM CTZ and 1.8–94 µM RLuc8 in the RLuc8-varying experiment, and 50 µM RLuc8 and 24 µM CTZ in the single turnover experiment.

*Product fluorescence stopped-flow kinetic experiments*

Transient kinetic traces upon the coelenteramide (CEI) product binding into the RLuc8 enzyme active site were monitored after rapidly mixing the components using the Stopped-Flow SFM 3000 mixing system equipped with a xenon arc lamp light source and the MOS-500 spectrometer (BioLogic, France).

The measurement was initiated by mixing 75 µL of the enzyme solution with 75 µL of the CEI solution with a total flow rate of 13 mL.min$^{-1}$ and then monitored by the change of the product fluorescence upon binding using the 375 nm long-pass emission filter after the excitation at 330 nm. The experiment was performed in 100 mM potassium phosphate buffer pH 7.5 at 15 °C and each kinetic trace was collected in 6 consecutive technical replicates and then averaged. The resulting concentration of CEI after mixing was 10 µM while the concentration of RLuc8 varied from 1.2 to 342 µM to generate a concentration series.

*Anaerobic tryptophan and substrate fluorescence stopped-flow kinetic experiments*

100 mM potassium phosphate buffer pH 7.5 and water solutions were incubated in the anaerobic glovebox Belle MR2 (Belle Technology, UK) overnight while stirring to remove dissolved oxygen. The glovebox was filled with ultra-high purity (UHP) nitrogen atmosphere containing a residual oxygen concentration of 2.2 ppm, as determined by the oxygen meter O2M-3 (Belle Technology, UK). A bottle with lyophilized evacuated (oxygen-free) RLuc8 protein was put into the anaerobic glovebox as well, it was dissolved with anaerobic water and incubated for at least one hour to remove possible traces of dissolved oxygen. Finally, a nitrogen-flushed aliquot of CTZ dissolved in pure ethanol was put into the glovebox, incubated for approximately 20 minutes to remove dissolved oxygen, and then closed to prevent further evaporation of the sample. CTZ stock solution and reconstituted RLuc8 solution were then diluted with the anaerobic 100 mM potassium phosphate buffer pH 7.5 to the desired concentrations to generate the initial samples for the stopped-flow experiments. The solutions were subsequently loaded into the Stopped-Flow instrument KinetAsyst SF-61DX2 (TgK Scientific, UK) that was placed in the same anaerobic glovebox Belle MR2 (Belle Technology, UK). The experiment was initiated by rapidly mixing the components in a 1:1 ratio and then monitored in two modes using two different sources of the optical signal: (i) native tryptophan fluorescence and (ii) native substrate fluorescence. Quenching of the tryptophan fluorescence, caused by the CTZ substrate binding in the active site, was monitored using the 313–430 nm band-pass emission filter after the excitation at 295 nm. The increase of the substrate fluorescence, caused by binding into the rigidifying hydrophobic active site of the enzyme, was monitored using the 445 nm long-pass emission filter after the excitation at 415 nm. The experiment was performed in 100 mM potassium phosphate buffer pH 7.5 at either 25 °C or 7 °C and each kinetic trace was collected in 6 consecutive technical replicates and then averaged. The resulting concentration of RLuc8 after mixing was 2 µM while the concentration of CTZ varied from 0.4 to 74 µM to generate a concentration series.

*Time-resolved spectral absorbance stopped-flow kinetic experiments*

Transient kinetic traces upon the coelenterazine (CTZ) substrate conversion/binding by the RLuc8 enzyme were monitored after rapidly mixing the components using the Stopped-Flow instrument KinetAsyst SF-61DX2 (TgK Scientific, UK). The measurement was initiated by mixing the enzyme and the CTZ substrate solution in a 1:1 ratio and then monitored by the change of absorbance spectra over time using the KinetaScan CCD Detector (TgK Scientific, UK). The measured spectral changes originated from (i) CTZ/CEI (un)binding by the enzyme and (ii) CTZ-to-CEI conversion. The experiment was performed in 100 mM potassium phosphate buffer pH 7.5 at either 7 °C or 37 °C in two different experimental setups: (i) standard aerobic mixing monitoring the whole catalytic cycle; (ii) anaerobic mixing (details on anaerobization of the samples provided in the previous section) to dissect only the initial kinetic phases of the substrate binding with no follow-up conversion. Each kinetic trace was collected in 6 consecutive technical replicates and then averaged. The resulting concentrations after mixing were 74 µM RLuc8 and 64 µM CTZ in the aerobic experiments and 25 µM RLuc8 and 64 µM CTZ in the anaerobic experiment.

*Temperature-dependent rapid-mixing microfluidic chip kinetic experiments*

Transient kinetic traces upon the coelenterazine (CTZ) substrate binding by luciferase enzymes (Anc$^{HLD\text{-}RLuc}$, AncFT, AncFT-L14, and RLuc8) at different temperatures were collected after rapidly mixing the components using the advanced continuous-flow microfluidic chip allowing high-throughput data collection with minimal sample consumption and rapid heat-transfer in the setup described and published previously(*11*). The measurement was initiated by mixing the luciferase enzyme solution with the CTZ solution at the total flow rate of 2.1 µL.min$^{-1}$ and then monitored by the change of the substrate fluorescence upon binding using the 488 nm long-pass emission filter after the excitation at 405 nm. The experiment was performed in 100 mM potassium phosphate buffer pH 7.5 at temperatures varying from 9 to 22 °C for AncFT, AncFT-L14, and RLuc8 and from 22 to 49 °C for Anc$^{HLD-RLuc}$. The resulting concentration of CTZ after mixing was kept constant at 30 µM while the concentrations of the luciferase enzymes varied from 7 to 67 µM for Anc$^{HLD-RLuc}$, from 10 to 94 µM for AncFT, from 6 to 57 µM for AncFT-L14, and from 7 to 76 µM for RLuc8.

*Global numerical analysis of transient kinetic data*

All the transient kinetic curves obtained by the methods described in the corresponding sections above were directly analyzed globally by numerical fitting using KinTek Explorer 10 (KinTek Corporation, USA). The software allows for the input of a given kinetic model via a simple text description, and the program then derives the differential equations needed for numerical integration automatically. Numerical integration of rate equations searching a set of kinetic parameters that produce a minimum $x^2$ value was performed using the Bulirsch–Stoer algorithm with adaptive step size, and nonlinear regression to fit data was based on the Levenberg–Marquardt method. Residuals were normalized by sigma value for each data point. The standard error (S.E.) was calculated from the covariance matrix during nonlinear regression. In addition to S.E. values, a more rigorous analysis of the variation of the kinetic parameters was accomplished by confidence contour analysis using FitSpace Explorer (KinTek Corporation, USA). In these analyses, the lower and upper limits for each parameter were derived from the confidence contour obtained from setting $x^2$ threshold at 0.95. During the analysis procedure, multiple kinetic schemes were tested until the minimal pathway accurately accounting for all the collected data was identified together with the corresponding rate constants and activation energies. The whole process of data fitting from starting kinetic curves and defined observables up to the final kinetic scheme is described more in detail in **Supplementary Note 1**.

*Co-crystallization assays*

Crystallization experiments were done at 20 °C using the hanging-drop vapor-diffusion method in EasyXtal 15-well plates (Qiagen, Germany) with drops equilibrated against 500 µL of reservoir solution. For co-crystallization, AncFT-L14 enzyme concentrated to ~9 mg.mL$^{-1}$ was mixed with azaCTZ(*3*) in a 1:2 molar enzyme-ligand ratio. Crystals were obtained after mixing 1 µL of enzyme-ligand mixture with 1 µL of the precipitant solution consisting of 0.1 M SPG (succinic acid, sodium phosphate monobasic monohydrate, and glycine) buffer pH 9 and 25 % PEG 1500. The crystals were harvested after 5 to 7 days of incubation. All crystals were fished out, cryo-protected in the corresponding reservoir solution supplemented with 20 % glycerol, and cryo-cooled in liquid nitrogen for X-ray data collection.

*Diffraction data collection and data processing*

X-ray data were collected at PXIII beamline at SLS Synchrotron (Villigen, Switzerland) at the wavelength of 0.999 Å using a Pilatus 2M-F detector. The data were processed using XDS(*12*), and Aimless(*13*) was used for data merging. Initial phases were solved by molecular replacement using Phaser(*14*) implemented in Phenix(*15*) with AncFT (PDB ID: 7QXR)(*3*) employed as a search model. The refinement was carried out in cycles of automated refinement in phenix.refine program(*15*) and manual model building in Coot(*16*). The final models were validated using tools provided by Coot(*16*) and MolProbity(*17*). Visualizations of structural data were created using PyMOL 2.0 (Schrödinger LLC, USA). Structural superposition was carried out using the secondary structure matching (SSM) superimpose tool in the Coot(*16*). Atomic coordinates and structure factors of the enzyme-ligand complex were deposited in the Protein Data Bank (www.wwpdb.org)(*18*) under the PDB code 7OME.

*Preparation of ligands and enzyme structures for in silico simulations*

The structures of CTZ (substrate) and CEI (product) were prepared using Avogadro 1.2.0 software(*19*): the multiplicity of the bonds was edited to match the keto forms, all missing hydrogens were added, and the structures were minimized by the steepest descent algorithm in the Auto Optimize tool of Avogadro, using the Universal Force Field (UFF). For the molecular docking, the restrained electrostatic potential (RESP) charges of the ligands were derived by the RESP ESP charge Derive (R.E.D.) Server Development 2.0(*20*). Next, the AutoDock atom types were added, and PDBQT files were generated by MGLTools(*21*, *22*). For the molecular dynamics simulations, the *antechamber* module of AmberTools16(*23*) was used to calculate the charges for the ligands, add the atom types of the Amber force field, and compile them in a PREPI parameters file. Also, the *parmchk2* tool from AmberTools16 was used to create additional FRCMOD parameter files to compensate for any missing parameters. The crystal structures of RLuc8, AncFT-L14, and Anc$^{HLD\text{-}RLuc}$ (chains A of PDB IDs 2PSF, 7OME, and 6G75, respectively) were downloaded from the RCSB Protein Data Bank(*18*) and stripped of all HETATM records (non-protein atoms). The structures were aligned to chain A of 2PSF in PyMOL 2.5.4 (Schrödinger LLC, USA) and protonated with H++ web server v. 4.0(*24*, *25*), using pH = 7.5, salinity = 0.1 M, internal dielectric = 10, and external dielectric = 80 as parameters. For the subsequent docking, AutoDock atom types and Gasteiger charges were added to the enzymes by MGLTools(*21*, *22*), and the corresponding PDBQT files were generated.

*Molecular docking*

The AutoDock Vina 1.1.2(*26*) software was used for molecular docking. For docking to the active site, the docking grid was specified as an x = 33.00 Å, y = 33.38 Å, z = 32.62 Å sized box, with a center in x = 47.83, y = 21.98, z = 12.41 Å, covering the catalytic pocket and access tunnels. The flag --exhaustiveness = 100 was used to sample the possible conformational space thoroughly. The number of output conformations of the docked ligand was set to 10. The results were analyzed in PyMOL 2.5.4 (Schrödinger LLC, USA).

*System preparation and equilibration for ASMD*

RLuc8, AncFT-L14, and Anc$^{HLD\text{-}RLuc}$ structures were prepared as described above. Then, the crystallographic water molecules were added back into the systems, except those overlapping with the protein or ligand. Next, histidine residues were renamed according to their protonation state (HID - Nδ protonated, HIE – Nε protonated, HIP – both Nδ and Nε protonated). The ligand was prepared as described in the corresponding section above.

Twelve systems were prepared for the CTZ binding analysis: four different CTZ poses were placed at the tunnel mouth of each studied enzyme using PyMOL (Schrödinger LLC, USA). The *tLEaP* module of AmberTools16(*23*) was used to neutralize the systems with $Cl^-$ and $Na^+$ ions, import the ff14SB force field(*27*) to describe the protein and the PREPI file parameters to describe the ligand, add a truncated octahedral box of TIP3P water molecules(*28*) to the distance of 10 Å from any atom in the system, and generate the topology file, coordinate file, and PDB file.

The system equilibration was carried out with the *PMEMD.CUDA*(*29–31*) module of Amber 16(*23*). Five minimization steps and twelve steps of equilibration dynamics were performed. The first four minimization steps comprised 2,500 cycles of the steepest descent algorithm followed by 7,500 cycles of the conjugate gradient algorithm, while harmonic restraints gradually decreased. The restraints were applied as follows: 500 kcal.mol$^{-1}$.Å$^{-2}$ on all heavy atoms of the protein and ligand, and then 500, 125, and 25 kcal.mol$^{-1}$.Å$^{-2}$ on protein backbone atoms and ligand heavy atoms. The fifth step comprised 5,000 cycles of the steepest descent algorithm followed by 15,000 cycles of the conjugate gradient algorithm without restraint.

The equilibration MD simulations consisted of twelve steps: (i) The first step involved 20 ps of gradual heating from 0 to 298 K at constant volume using Langevin dynamics, with harmonic restraints of 200 kcal.mol$^{-1}$.Å$^{-2}$ on all heavy atoms of the protein and ligand, (ii) ten steps of 400 ps equilibration Langevin dynamics each at a constant temperature of 298 K and a constant pressure of 1 bar with decreasing harmonic restraints of 150, 100, 75, 50, 25, 15, 10, 5, 1, and 0.5 kcal.mol$^{-1}$.Å$^{-2}$ on protein backbone and ligand heavy atoms, and (iii) the last step involving 400 ps of equilibration dynamics at a constant temperature of 298 K and a constant pressure of 1 bar with no restraint. The simulations employed periodic boundary conditions based on the particle mesh Ewald method(*32*) for treatment of the long-range interactions beyond the 10 Å cut-off, the SHAKE algorithm(*33*) to constrain the bonds that involve hydrogen atoms, the Berendsen barostat(*34*) at 1 bar, and the Langevin temperature equilibration using a collision frequency of 1 ps$^{-1}$.

After the equilibration, the number of $Cl^-$ and $Na^+$ ions needed to reach 0.1 M salinity was calculated using the average volume of the system in the last equilibration step. The whole process was repeated, from the *tLEaP* step, to correct the number of the added ions.

*Adaptive steered molecular dynamics*

The CTZ binding trajectories were calculated with adaptive steered molecular dynamics (ASMD). The ASMD method applies constant external force on two atoms in the simulated systems. By pulling two atoms together, we simulated ligand binding. During ASMD, several parallel simulations were started from the same state. The simulation ran in stages where the distance between the selected atoms changed by 2 Å per stage. At the end of each stage, the parallel simulations were collected, analyzed, and the Jarzynski average(*35*, *36*) was calculated throughout the stage. The trajectory with its work value closest to the Jarzynski average was selected, and the state at the end of this trajectory was used as the starting point for the next stage. We used the default values for setting up ASMD from the respective Amber tutorial and the ASMD publication(*37*). The simulations were run with 25 parallel MDs, steered by 2 Å stages of distance increments, with a velocity of 10 $Å.ns^{-1}$ and a force of 7.2 N. The rest of the MD settings were set as in the last equilibration step. To simulate CTZ binding, we selected the N1-nitrogen of CTZ and the Cγ atom of D120 residue as steering atoms (D120 in RLuc8 corresponds to D118 in the ancestral variants). The initial distance between the steering atoms in the equilibrated systems was measured using PyMOL (Schrödinger LLC, USA). The selected steering atoms were pulled together to the distance of 4.4 Å, as observed in the crystal structure PDB ID 7QXR (azaCTZ-bound AncFT luciferase).

*System preparation and equilibration for adaptive sampling*

The adaptive sampling method was used to simulate product release from the studied enzymes with a thorough sampling of the unbinding process. The systems for adaptive sampling were built using the crystal structures of RLuc8, AncFT-L14, and $Anc^{HLD\text{-}RLuc}$. Three systems were generated with CEI placed in the active site, as seen in PDB ID 7QXQ (AncFT + CEI). An additional system was prepared for AncHLD-RLuc with a flipped CEI, as observed in the adaptive steered MD simulation of CTZ binding.
The following steps were performed with the High Throughput Molecular Dynamics (HTMD)(*38*) scripts. The protein structures were protonated with PROPKA 2.0(*39*) at pH 7.5. Each system was solvated in a cubic water box of TIP3P(*28*) water molecules with the edges at least 10 Å away from the protein by the *solvate* module of HTMD. $Cl^-$ and $Na^+$ ions were added to neutralize the charge of the protein and get a final salt concentration of 0.1 M. The topology of the system was built using *amber.build* module of HTMD, with the ff14SB(*27*) Amber force field and the previously compiled PREPI and FRCMOD parameter files for the ligands. The systems were equilibrated using the *equilibration_v2* module of HTMD3(*38*). The system was first minimized using a conjugate-gradient method for 500 steps. Then the system was heated to 298 K and minimized as follows: (i) 500 steps (2 ps) of NVT thermalization with the Berendsen barostat with 1 $kcal.mol^{-1}.Å^{-2}$ constraints on all heavy atoms of the protein, (ii) 1,250,000 steps (5 ns) of NPT equilibration with Langevin thermostat and same constraints, and (iii) 1,250,000 steps (5 ns) of NPT equilibration with the Langevin thermostat without any constraints. During the equilibration simulations, holonomic constraints were applied on all hydrogen-heavy atom bond terms, and the mass of the hydrogen atoms was scaled with factor 4, enabling time steps of 4 fs(*40*–*43*). The simulations employed periodic boundary conditions, using the particle mesh Ewald method to treat interactions beyond the 9 Å cut-off. The 1-4 electrostatic interactions were scaled with a factor of 0.8333, and the smoothing and switching of van der Waals interaction was performed for a cut-off of 7.5 Å(*42*).

*Adaptive sampling*

HTMD3(*38*) was used to perform adaptive sampling of the conformations of the systems. Production MD runs of 50 ns were started using the equilibrated systems, employing the same settings as in the last equilibration step. The trajectories were saved every 0.1 ns. Adaptive sampling was performed using the distance of the N1-nitrogen of CEI and the Cγ atom of catalytic residue D120 as the adaptive metric (residue D120 in RLuc corresponds to D118 in the ancestral variants), and time-lagged independent component analysis (tICA)(*44*) projection in 1 dimension. 20 epochs of 10 parallel MDs were performed, corresponding to a cumulative time of 10 µs per system.

*Markov state model construction*

The simulations were made into a simulation list using HTMD3(*38*); the water and ions were filtered out, and unsuccessful simulations shorter than 50 ns were omitted. Such filtered trajectories were combined, which resulted in 10 µs of cumulative simulation time for each of the five systems.

CEI release was studied based on the distance of the N1-nitrogen of CEI and the Cγ atom of the D120 residue (D120 in RLuc corresponds to D118 in the ancestral variants). The data were clustered using the MiniBatchKmeans algorithm to 1,000 clusters. Markov state models (MSM) with three macrostates were constructed at a lag time of 17 ns for each system with crystal-like CEI and at 15 ns for the flipped CEI unbinding. The three macrostates were labeled based on the CEI location as 'bound', 'tunnel', and 'unbound'.

The opening of the access tunnel (active site opening caused by the conformational change of the α4 helix) was studied based on the distance of the Cα atoms of I159/V185 residues in RLuc8, I157/L183 in AncFT-L14, and A158/L184 in $Anc^{HLD\text{-}RLuc}$. The data were clustered using the MiniBatchKmeans algorithm to 1,000 clusters. MSMs with two macrostates for each system were constructed at a lag time of 17 ns for RLuc8, 20 ns for AncFT-L14 and $Anc^{HLD\text{-}RLuc}$ with crystal-like CEI, and 15 ns for $Anc^{HLD\text{-}RLuc}$ with flipped CEI.

The Chapman-Kolmogorov test was performed to ensure the models are Markovian and describe the data well (**Figure S14** and **S15**). The states were visualized in VMD 1.9.3(*45*), and the statistics of the ligand-active site distance were calculated (mean distance, standard error S.E., minimum, and maximum). The trajectory was saved for each model and visualized in PyMOL (Schrödinger LLC, USA).

*Kinetics analysis of molecular dynamics simulations*

Kinetic values (MFPT on/off, $k_{on}$, $k_{off}$, $k_{off}/k_{on}$, $\Delta G^0_{eq}$, and $K_d$) were calculated by the *kinetics* module of HTMD3(*38*) between the source ('unbound' state) and sink ('bound' state). Also, the equilibrium probability of each macrostate (population of a specific conformation observed during the simulation) was calculated and visualized. Finally, bootstrapping of the kinetics calculation was performed, using randomly selected 80 % of the data, run 100 times. The kinetic values were then averaged, and the standard errors were determined.

*Binding energy calculation using MM/GBSA*

The molecular mechanics/generalized Born surface area (MM/GBSA)(*46*, *47*) method was applied to calculate CEI binding free energy ($\Delta G_{bind}$) in the enzymes and the respective residue-by-residue interactions. Ante-MMPBSA.py module of AmberTools 16(*23*) was used to remove the solvent and ions from the original topology files of each system, define the Born radii as mbondi3, and generate the corresponding topology files for the complex, receptor, and ligand. The $\Delta G_{bind}$ of the ligand in each structure was calculated with the MMPBSA.py program(*46*). The generalized Born method was used (*&gb* namelist) with the implicit generalized Born solvent model (*igb=8*) and salt concentration 0.1 M (*saltcon=0.1*). The solvent-accessible surface area was computed with an LCPO algorithm(*48*). Decomposition of the interactions (*&decomp* namelist) was generated per residue (*idecomp=2*), with discrimination of all types of energy contributions (*dec_verbose=0*). The MM/GBSA energies were calculated on 1,000 snapshots of each enzyme's 'bound' and 'tunnel' macrostates.
